# Dynamic Predictive Coding: A Model of Hierarchical Sequence Learning and Prediction in the Neocortex

**DOI:** 10.1101/2022.06.23.497415

**Authors:** Linxing Preston Jiang, Rajesh P. N. Rao

## Abstract

We introduce dynamic predictive coding, a hierarchical model of spatiotemporal prediction and sequence learning in the neocortex. The model assumes that higher cortical levels modulate the temporal dynamics of lower levels, correcting their predictions of dynamics using prediction errors. As a result, lower levels form representations that encode sequences at shorter timescales (e.g., a single step) while higher levels form representations that encode sequences at longer timescales (e.g., an entire sequence). We tested this model using a two-level neural network, where the top-down modulation creates low-dimensional combinations of a set of learned temporal dynamics to explain input sequences. When trained on natural videos, the lower-level model neurons developed spacetime receptive fields similar to those of simple cells in the primary visual cortex while the higher-level responses spanned longer timescales, mimicking temporal response hierarchies in the cortex. Additionally, the network’s hierarchical sequence representation exhibited both predictive and postdictive effects resembling those observed in visual motion processing in humans (e.g., in the flash-lag illusion). When coupled with an associative memory emulating the role of the hippocampus, the model allowed episodic memories to be stored and retrieved, supporting cue-triggered recall of an input sequence similar to activity recall in the visual cortex. When extended to three hierarchical levels, the model learned progressively more abstract temporal representations along the hierarchy. Taken together, our results suggest that cortical processing and learning of sequences can be interpreted as dynamic predictive coding based on a hierarchical spatiotemporal generative model of the visual world.

**Author Summary:** The brain is adept at predicting stimuli and events at multiple timescales. How do the neuronal networks in the brain achieve this remarkable capability? We propose that the neocortex employs dynamic predictive coding to learn hierarchical spatiotemporal representations. Using computer simulations, we show that when exposed to natural videos, a hierarchical neural network that minimizes prediction errors develops stable and longer timescale responses at the higher level; lower-level neurons learn space-time receptive fields similar to the receptive fields of primary visual cortical cells. The same network also exhibits several effects in visual motion processing and supports cue-triggered activity recall. Our results provide a new framework for understanding the genesis of temporal response hierarchies and activity recall in the neocortex.

## 1 Introduction

The ability to predict future stimuli and event outcomes is critical for perceiving and interacting with a highly dynamic world. At the neural circuit level, predictions could compensate for neural transmission delays and engage with the world in real-time. At the cognitive level, planning a sequence of actions to achieve a desired goal relies on predictions of the sensory consequences of motor commands. These abilities are predicated on two requirements: (a) the brain must infer the dynamics of sensory stimuli to make spatiotemporal predictions based on an internal model of the world, and (b) the brain’s temporal representations must span different timescales to support predictions over both short and long horizons.

Many experimental studies have provided evidence for such computations. Predictive representations of upcoming stimuli have been found in various open and closed-loop paradigms where animals developed experience-dependent visual and auditory expectations [1–5]. Other empirical evidence suggests that cortical representations exhibit a hierarchy of timescales and an increase in stability from lower-order to higher-order areas across both sensory and cognitive regions [6–9]. We asked the question: could such phenomena be explained by the neocortex learning a spatiotemporal generative model based on a temporal hierarchy of representations?

Predictive coding provides a unifying framework for understanding perception and prediction in terms of learning hierarchical generative models of the environment [10–14]. Here, we present dynamic predictive coding (DPC), a new predictive coding model for learning hierarchical temporal representations. The central idea of our proposal is that our perceptual system learns temporally abstracted representations that encode entire sequences rather than single points at any given time. Specifically, DPC assumes that higher-level model neurons modulate the *transition dynamics* of lower-level networks, building on the computational concept of *hypernetworks* [15]. Hypernetworks are neural networks that generate the parameters (synaptic weights) for another neural network. However, generating an entire set of high-dimensional synaptic weights is not neurally plausible. Instead, DPC models the transition dynamics at a lower level of a hierarchy using a small set of modulation weights for a group of learned transition matrices. These weights implement “top-down” gain modulation of the lower-level synapses [16, 17] and are predicted by the higher level through a feedback network (a hypernetwork) connecting the higher to the lower level. Compared to previous normative models of video processing that either do not learn the temporal dynamics between images [18–22] or presume a fixed temporal hierarchy [23, 24] (see Discussion), the DPC model offers a neural implementation of spatiotemporal prediction that learns the transition dynamics of the input and adapts its hierarchical temporal representation to the intrinsic timescales of the data.

We tested the DPC model using a two-level neural network trained on natural and artificial image sequences to minimize spatiotemporal prediction errors. After training, the lower-level neurons developed space-time receptive fields similar to those found in simple cells in the primary visual cortex (V1) [25]. Neurons in the second level learned to capture input dynamics on a longer timescale and their responses exhibited greater stability compared to responses in the first level, similar to the temporal response hierarchies observed in the cortex [6–9]. We further show that the learned sequence representations in the network can explain both predictive and postdictive effects seen in visual processing [26–29], reproducing several aspects of the flash-lag illusion [26, 30, 31]. When linked to an associative memory mimicking the role of the hippocampus, the network allowed storage of episodic memories and exhibited cue-triggered activity recall after repeated exposure to a fixed input sequence, an effect previously reported in rodents [1], human V1 [32–34] and monkey V4 [35]. Lastly, when extended to three levels, the top-level neurons learned to encode the transition dynamics of the second-level states, which in turn encoded the transition dynamics of the first-level states, thereby yielding a hierarchical temporal representation of input image sequences. Together, these results support the hypothesis that the neocortex uses dynamic predictive coding based on a hierarchical spatiotemporal generative model to learn and interpret input sequences at multiple levels of temporal abstractions. Some of the results presented herein appeared previously in a conference proceedings [36].

## 2 Results

### Dynamic predictive coding

The DPC model assumes that spatiotemporal inputs are generated by a hierarchical generative model (Figure 1a) (see also [37]). We describe here a two-level hierarchical model (see Discussion for the possibility of extending the model to more levels). The lower level of the model follows the traditional predictive coding model in generating images using a set of spatial filters **U** and a latent state vector **r**_*t*_, which is sparse [38], for each time step *t*: **I**_*t*_ = **Ur**_*t*_ + **n** where **n** is zero mean Gaussian white noise. The temporal dynamics of the state **r**_*t*_ is modeled using *K* learnable transition matrices 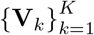 which can be linearly combined using a set of “modulation” weights given by a *K*-dimensional vector **w**. This vector of weights is generated by the higher-level state vector **r**^*h*^ using a function ℋ (Figure 1(b)), implemented as a neural network (a “hypernetwork” [15] – see Supplementary Information):

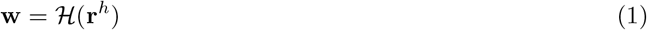

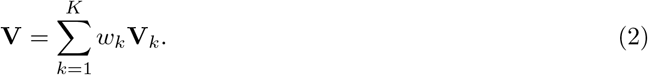

Here, *w*_*k*_ is the *k*th component of the vector **w**. The lower-level state vector at time *t* + 1 is generated as **r**_*t*+1_ = ReLU(**Vr**_*t*_) + **m** where **m** is zero mean Gaussian white noise. Note that this is one particular parameterization for top-down modulation of the lower-level transition dynamics, with the hypernetwork formulation allowing other types of parameterizations (see Supplementary Information).

**Figure 1:**
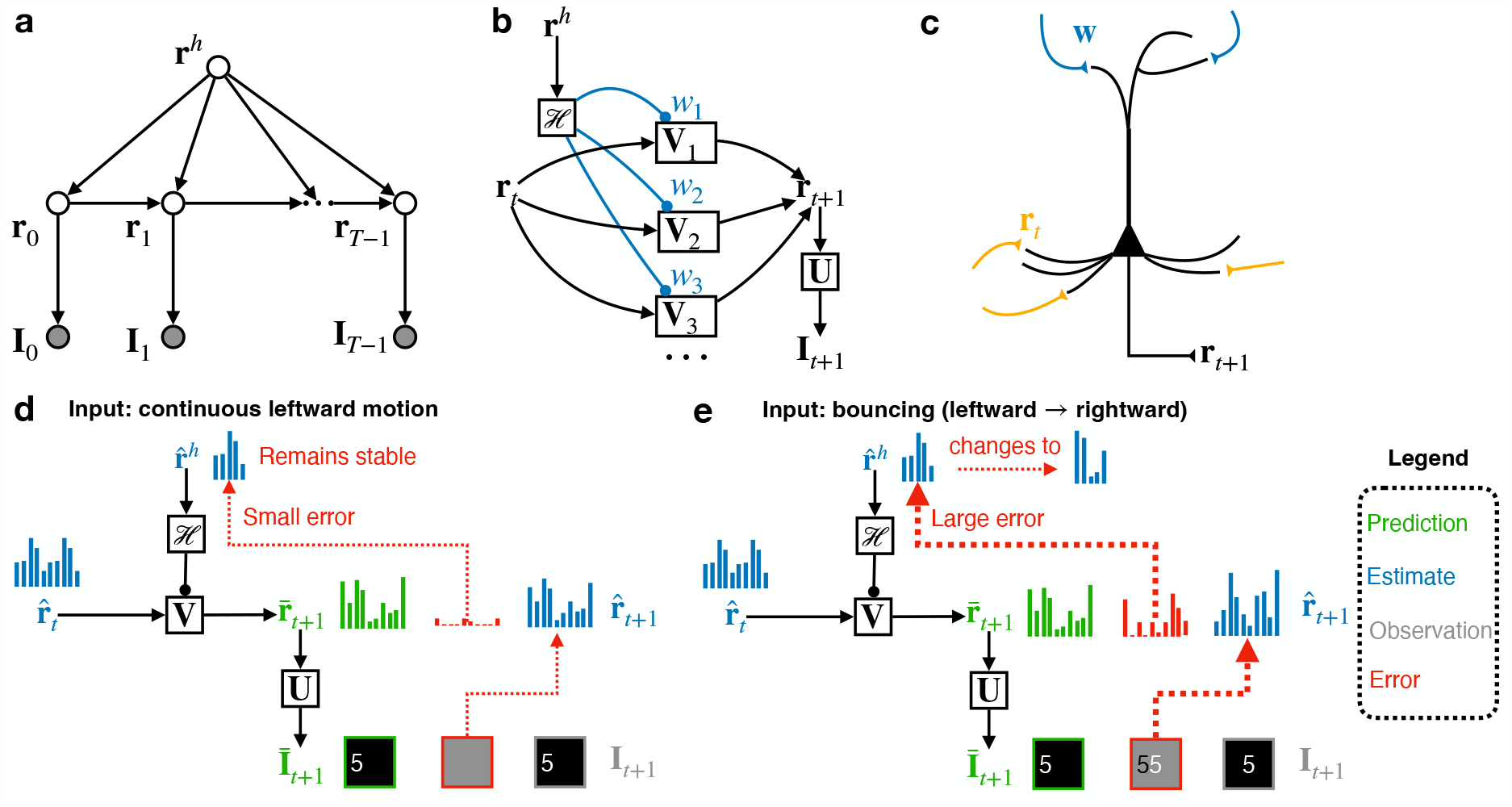
Dynamic predictive coding. **(a)** Generative model for dynamic predictive coding. **(b)** Parameterization of the model. The higher-level state modulates the lower-level transition matrices through a top-down network (“hypernetwork”) ℋ. **(c)** A possible neural implementation of the generative model using cortical pyramidal neurons. Pyramidal neurons receive the top-down embedding vector input via synapses at apical dendrites and the current recurrent state vector via basal dendrites, and produce as their output the next state vector. **(d)** Schematic depiction of an inference step when the dynamics at the lower level is stable. The higher-level state remains stable due to minimal prediction errors. **(e)** Depiction of an inference step when the lower-level dynamics changes. The resulting large prediction errors drive updates to the higher-level state to account for the new lower-level dynamics.

The generative model in Figure 1(b) can be implemented in a hierarchical neural network: the higherlevel state **r**^*h*^, represented by higher-level neurons, generates a top-down modulation **w** via a top-down feedback neural network *ℋ*, and this top-down input **w** influences the groups of lower-level neurons representing **V**_*i*_ through gain modulation [16, 17] (see Supplementary Information for details). We propose that such a computation could be implemented by cortical pyramidal neurons receiving topdown modulation via their apical dendrites (through gain control [17, 39]) and the recurrent state **r**_*t*_ (and input prediction errors) via their basal dendrites, and integrating these to predict the next state (Figure 1(c)).

When an input sequence is presented, the model employs a Bayesian filtering approach to perform online inference on the latent vectors [40] by minimizing a loss function that includes prediction errors and penalties from prior distributions over the latent variables (see Methods). Given the model’s estimates 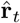 and 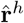 at time *t*, the estimate 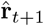 of **r** at time *t*+1 is computed by gradient descent to minimize the sum of the input prediction error 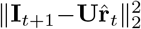 and the temporal state prediction error 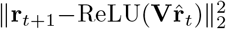 plus a sparseness penalty. Similarly, the second level estimates 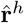 is updated using the temporal prediction error plus a prior-related penalty. The model’s parameters are learned by minimizing the same prediction errors across all time steps and input sequences, further reducing the errors not accounted for by the inference process above for latent vectors (see Methods).

### Hierarchical predictive coding of natural videos

We implemented the DPC model described above using a two-level neural network where neural responses represent estimates of the latent state vectors and whose synaptic weights represent the spatial filters and transition parameters. We used *K* = 5 transition matrices for the first level (more matrices did not significantly improve performance – see Supplementary Figure S1). Perception in the DPC network corresponds to estimating the latent vectors by updating neural responses (through network dynamics) to minimize prediction errors via gradient descent (see Methods). Updating network parameters to further reduce prediction errors corresponds to learning (slow changes in synaptic weights through synaptic plasticity).

Figures 1(d) & 1(e) illustrate the inference process for both levels of the network. The network generates top-down and lateral predictions (green) using the current two-level state estimates (blue). If the input sequence is predicted well by the top-down-modulated transition matrix **V**, the higher-level response **r**^*h*^ remains stable due to small prediction errors (Figure 1(d)). When a non-smooth transition occurs in the input sequence, the resulting large prediction errors are sent to the higher level via feedforward connections (red arrows, Figure 1(e), driving changes in **r**^*h*^ to predict new dynamics for the lower level.

We trained the network on thousands of natural image sequences extracted from a video recorded by a person walking on a forest trail (frame size: 16 *×* 16 pixels, sequence length: 10 frames ( ∼0.35 seconds)). The frames were spatially and temporally whitened to simulate retinal and lateral geniculate nucleus (LGN) processing [38, 41]. The image sequences reserved for testing did not overlap in space or time with the training sequences. Figure 2(a) illustrates the inference process on an example natural image sequence by the network. The first row displays the ground truth input **I**_*t*_ for 10 time steps: each frame was shown sequentially to the model. The next row shows the model’s predictions 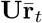 for each time step *t*, where 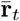 was predicted by the previous state estimate 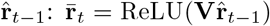. The prediction errors 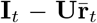 are shown in the third row. The prediction errors were the largest in the first two steps as the model inferred the spatial features and the transition dynamics from the initial inputs. The subsequent predictions were more accurate, resulting in minimized prediction errors. Finally, the last row shows the corrected estimates 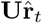 after 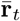 has been updated to 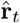 through prediction error minimization. Figure 2(b) shows the lower-(top) and higher-level (middle) neural responses to the natural video sequence in Figure 2(a). The bottom panel of Figure 2(b) shows the top-down dynamics modulation generated by the higher level.

**Figure 2:**
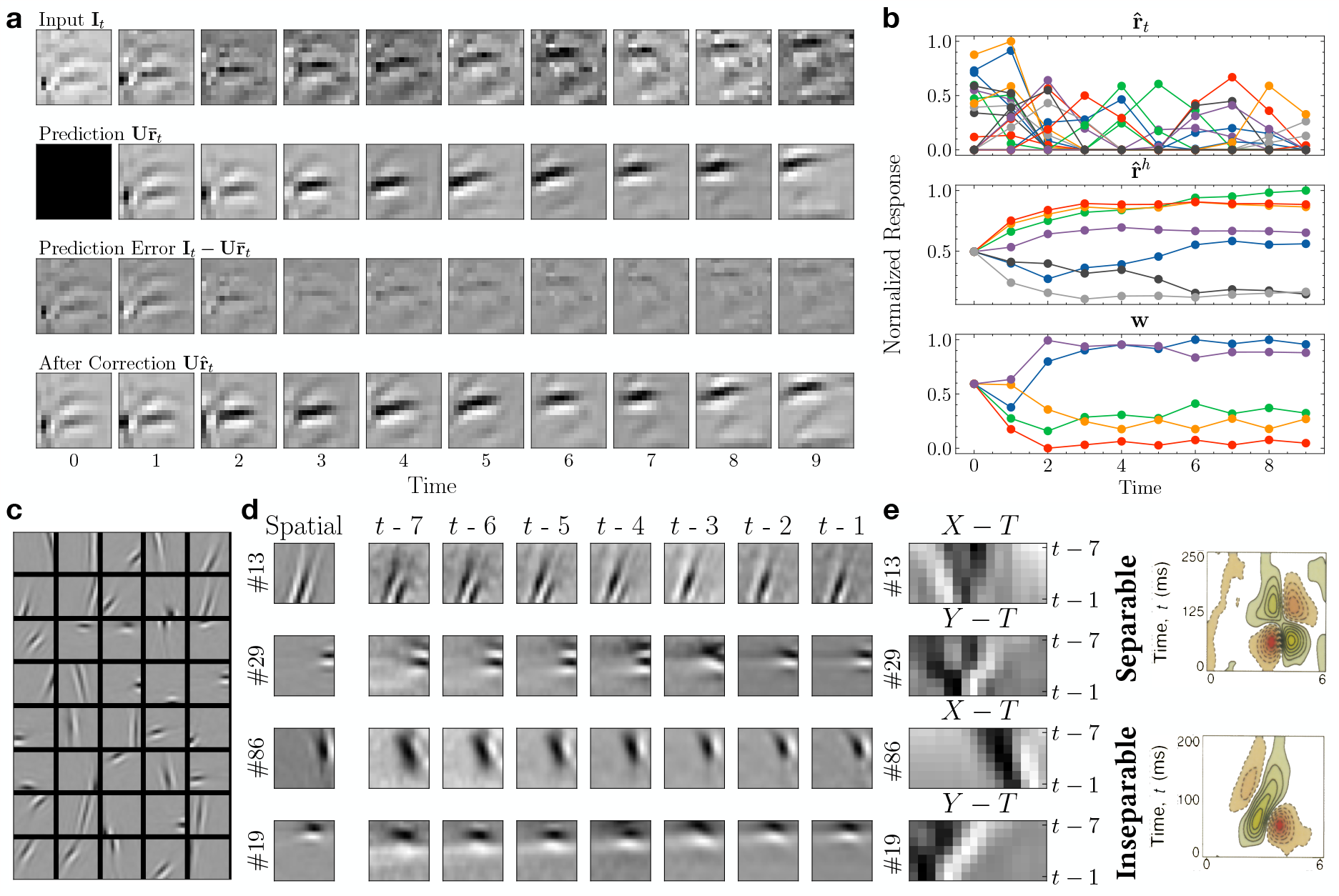
Predictive coding of natural videos and learned space-time receptive fields. **(a)** Inference on an example input image sequence of 10 frames. Top to bottom: Input sequence; model’s prediction of the current input from the previous step (the first step prediction being zero); prediction error (predicted input subtracted from the actual input); model’s final estimate of the current input after prediction error minimization. **(b)** The trained DPC network’s response to the natural image sequence in (a). Each plotted line represents the responses of a model neuron over 10 time steps. Top: responses of the 20 most active lower-level neurons (some colors are repeated); middle: responses of seven randomly chosen higher-level neurons; bottom: predicted transition dynamics (each line is the modulation weight for a basis transition matrix at the lower level). **(c)** 40 example spatial receptive fields (RFs) learned from natural videos. Each square tile is a column of **U** reshaped to a 16 *×* 16 image. **(d)** Space-Time RFs (STRFs) of four example lower-level neurons. First column: the spatial RFs of the example neurons. Next seven columns: the STRFs of the example neurons revealed by reverse correlation mapping. **(e)** Left panel: space-time plots of the example neurons in (d). Right panel: space-time plots of the RFs of two simple cells in the primary visual cortex of a cat (adapted from [25]).

We examined the learned spatial receptive fields (RFs) of the model neurons at the first level and qualitatively compared them with the spatial RFs of simple cells in the primary visual cortex (V1). A subset of the spatial filters (columns of **U**) learned by the model from our natural videos dataset are shown in Figure 2(c). These filters resemble oriented Gabor-like edge or bar detectors, similar to the localized, orientation-selective spatial RFs found in V1 simple cells [38, 42]. To measure the spatiotemporal receptive fields of the lower-level neurons, we ran a reverse correlation experiment [43, 44] with a continuous natural video clip ( ≈ 47 minutes) extracted from the same forest trail natural video used for training. This video was not shown to the model during either training or testing (see Methods). Figure 2(d) shows the spatiotemporal receptive fields for four example lower-level model neurons, computed by weighting input frames from the seven previous time steps **I**_*t*−7_, **I**_*t*−6_, …, **I**_*t*−1_ by the response 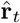 they caused at the current time step *t* (see Methods). The resulting average spatiotemporal receptive fields are shown as seven-image sequences labeled *t* − 7, *t* − 6, …, *t* − 1 (lasting ≈ 250 milliseconds in total). The first column labeled “Spatial” shows the spatial RFs of the example neurons.

To compute the space-time receptive fields (STRFs), we took the spatiotemporal *X* − *Y* − *T* receptive field cubes and collapsed either the *X* or *Y* dimension, depending on which axis had time-invariant responses. Figure 2(e) left panel shows the *X/Y − T* receptive fields of these example neurons. For comparison, Figure 2(e) right panel shows the STRFs of simple cells in the primary visual cortex (V1) of a cat (adapted from DeAngelis et al. [25]).

DeAngelis et al. [25] categorized the receptive fields of simple cells in V1 to be space-time separable (Figure 2(e) top row) and inseparable (Figure 2(e) bottom row). Space-time separable receptive fields maintain the spatial form of bright/dark-excitatory regions over time but switch their polarization: the space-time receptive field can thus be obtained by multiplying separate spatial and temporal receptive fields. Space-time inseparable receptive fields on the other hand exhibit bright/dark-excitatory regions that shift gradually over time, showing an orientation in the space-time domain. Neurons with spacetime inseparable receptive fields are direction-selective, responding to motion in only one direction. As seen in Figure 2(e) left pane, the neurons in the lower level of our network learned V1-like separable and inseparable STRFs, based on the principle of spatiotemporal prediction error minimization. To our knowledge, these results represent one of the first demonstrations of the emergence of both separable and inseparable STRFs in a recurrent network model by predictive coding of natural videos presented frame-by-frame. Previous demonstrations (e.g., [18, 20, 23]) have typically required chunks or all the frames of a video to be provided as a single input to a network, which is hard to justify biologically (see Discussion).

### Temporal hierarchy through prediction of dynamics

Next, we show that the two-level DPC network learned a hierarchical temporal representation of input videos. In our formulation of the model, a higher-level state vector predicts the *dynamics* of the lower-level states. This implies that the higher-level network neurons will have stable activation for input sequences with consistent dynamics (Figure 1(d)). When a change occurs in input dynamics, we expect the higher-level responses to switch to a different activation profile to minimize prediction errors (Figure 1(e)). We hypothesize that the different timescales of neural responses observed in the cortex [6–9] could be an emergent property of the cortex learning a similar hierarchical generative model.

We tested this hypothesis in our DPC network trained on natural videos. As seen in the inference example in Figure 2(b), the lower-level responses change rapidly as the stimulus moves (top panel). The higher-level responses (middle panel) and the predicted transition dynamics (right panel) were more stable after the initial adaptation to the motion. Since the stimulus continued to follow roughly the same dynamics (leftward motion) after the first two steps, the transition matrix predicted by the higher-level neurons continued to be accurate for the rest of the steps, leading to small prediction errors and few changes in the responses. Note that we did not enforce a longer time constant or smoothness constraint for **r**^*h*^ during inference – the longer timescale and more stable responses are entirely a result of the higher-level neurons learning to predict the lower-level dynamics under the proposed generative model.

To quantify this learned hierarchical temporal representation, we computed the autocorrelation of the lower- and higher-level responses to unseen natural videos and fitted an exponential decay function (see Methods). As Figure 3(a) shows, the autocorrelation of the higher-level responses **r**^*h*^ is greater than that of the lower-level response **r** and has a slower decay rate (exponential time constant for **r**^*h*^: 5.49 steps; for **r**: 2.18 steps). To factor out the effect of the natural video statistics, we computed the same autocorrelation using Gaussian white noise sequences. We found that the hierarchy of timescales still exists (**r**^*h*^: 3.11 steps; **r**: 0.18 steps). Figure 3(b) shows the autocorrelation of nonhuman primate neural responses from the medial-temporal (MT) area in the visual cortex (assumed to be lower in the processing stream) and lateral prefrontal cortex (LPFC) (assumed to be higher) for time periods preceding a motion task [6]. These neural responses show a difference in timescales qualitatively similar to the different response timescales in our hierarchical DPC network.

**Figure 3:**
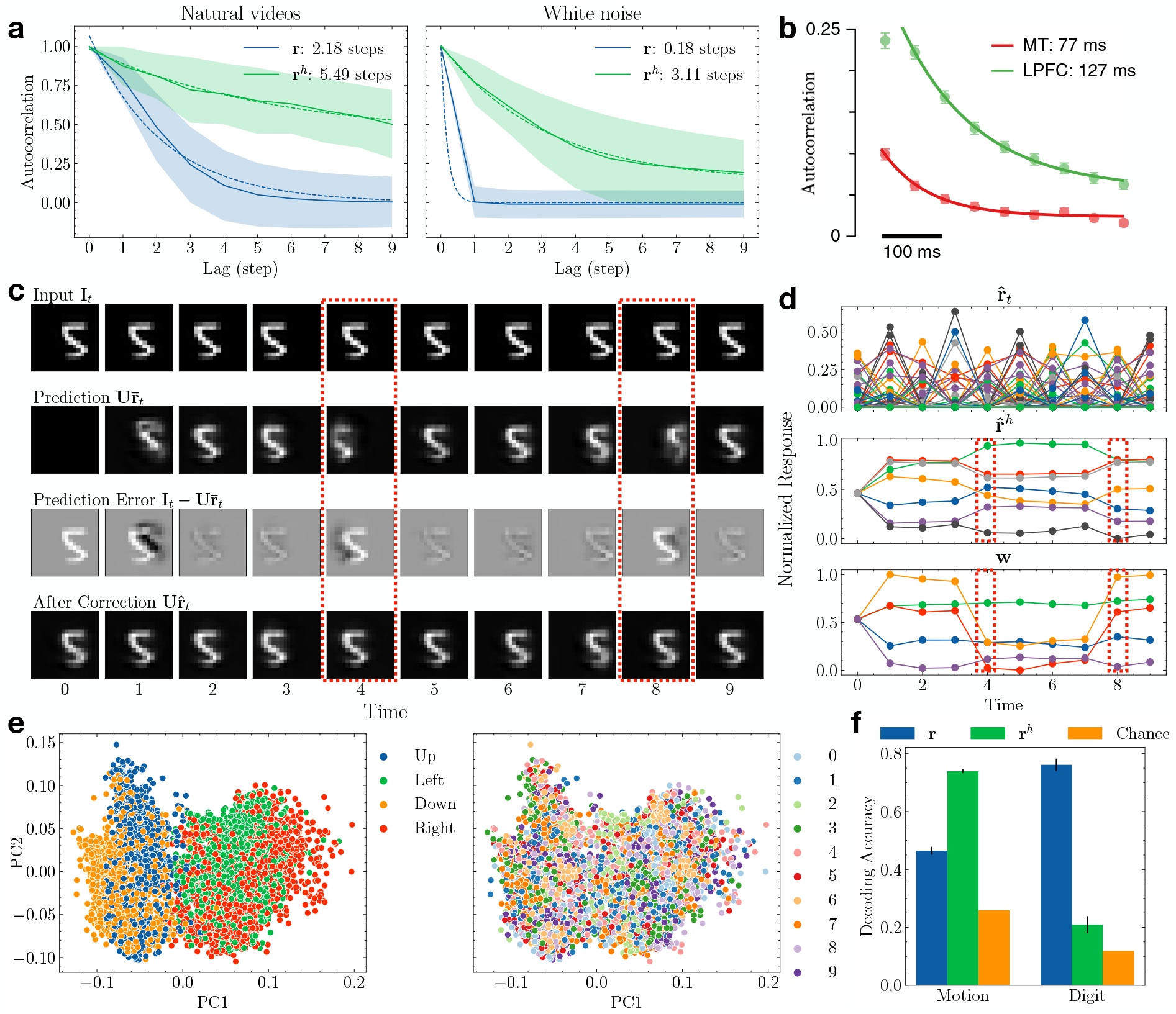
Hierarchical temporal representation with different timescales. **(a)** Autocorrelation of the lower- and higher-level responses in the trained network with natural videos. Shaded area denotes *±* 1 standard deviation. Dotted lines show fitted exponential decay functions. Left: response recorded during natural video stimuli; right: white noise stimuli. **(b)** Autocorrelation of the neural responses recorded from MT and LPFC of monkeys. Adapted from Murray et al. [6] **(c)** Inference for an example Moving MNIST sequence in a trained network. The red dashed boxes mark the time steps when the dynamics of the input changed. **(d)** The network’s responses to the input Moving MNIST sequence in (c). Note the changes in the higher-level responses after the input dynamics changed (red dashed boxes); this gradient-based change helps to minimize prediction errors. **(e)** Higher-level responses to the Moving MNIST sequences visualized in the 2D space of the first two principal components. Left: responses colored according to motion direction; right: responses colored according to digit identities. **(f)** Comparison of decoding performance for motion direction versus digit identity using lower- and higher-level neural responses. Error bars: *±* 1 standard deviation from 10-fold cross validation. Orange: chance accuracies.

To further understand the model’s ability to learn hierarchical temporal representations, we trained a DPC network on the Moving MNIST dataset [45]. Each image sequence in this dataset contains ten 18 *×* 18 pixel frames showing a single example of a handwritten digit (chosen from the original MNIST dataset) moving in a particular direction. The digit’s motion is limited to up, down, left, or right directions with a fixed speed. Figure 3(c) illustrates the trained network’s inference process on an example image sequence. Similar to the responses to the natural video sequence, the lower-level responses displayed fast changes while the higher-level responses spanned a longer timescale and showed greater stability (Figure 3(d)). Note that at time *t* = 4 and *t* = 8, the input dynamics changed as the digit “bounced” against the boundaries and started to move in the opposite motion (Figure 3(c) red dashed box). The higher-level neurons’ predictions resulted in large prediction errors at those times (Figure 3(c) third row). The prediction errors caused notable changes in the higher-level responses **r**^*h*^ (Figure 3(d) red dashed boxes). For the rest of the steps, **r**^*h*^ remained stable and generated accurate predictions of the stable dynamics.

Lastly, we confirmed that lower-level transition dynamics are indeed encoded in the higher-level responses.

We performed principal component analysis (PCA) on the higher-level responses **r**^*h*^ for the Moving MNIST sequences in the test set. Figure 3(e) visualizes these responses in the space of their first two principal components (PCs), colored by either the motion direction (left) or digit identities (right). The responses clearly formed clusters according to input motion direction but not digit identities. We then trained a support vector machine with radial basis function (RBF) kernel [46] to map **r** and **r**^*h*^ to motion directions and digit identities (Figure 3(f)). Using the higher-level responses, the classifier yielded 73.9% 10-fold cross-validated classification accuracy on the four motion directions (chance accuracy: 26.0%, computed as the number of majority labels in the test set divided by the total number of labels). Using the lower-level responses resulted in significantly less classification accuracy for motion direction (46.5%, *p* ≪ 0.001, *t*-test). In contrast, decoding accuracy for digit identity was significantly higher using the lower-level responses (76.1%) compared to using the higher-level responses (20.9%, *p* ≪ 0.001, *t*-test). These results show that due to the structure of its generative model, the DPC network learned to disentangle to a significant extent the motion information in an input video from image content (here, digit identity), yielding a factored representation of input image sequences.

### Predictive and postdictive effects in visual motion processing

The ability of the DPC model to encode entire sequences at the higher level (cf. the “timeline” model of perception [29]) leads to new normative and computational interpretations of visual motion phenomena such as the flash-lag illusion [26, 30, 31], explaining both predictive and postdictive effects [27, 29]. The flash-lag illusion refers to the phenomenon that a flashed, intermittent object is perceived to be “lagged” behind the percept of a continuously moving object even though the physical locations of the two objects are aligned or the same [30, 31]. Though this illusion is commonly attributed to the predictive nature of the perceptual system [30], Eagleman and Sejnowski [26] proposed a postdictive mechanism based on psychophysical results that the motion of the moving object *after* the flash can change the percept of events at the time of the flash. The potential interplay between prediction and postdiction in shaping perception was also studied by Hogendoorn et al. [27, 29]. The authors designed an interference paradigm with different reaction-speed tasks and showed that when the trajectory of the object unexpectedly reverses, predictive effects (extrapolation) are observed at short latencies but postdictive effects (interpolation) are observed at longer latencies (Figure 4(i)&(j)).

**Figure 4:**
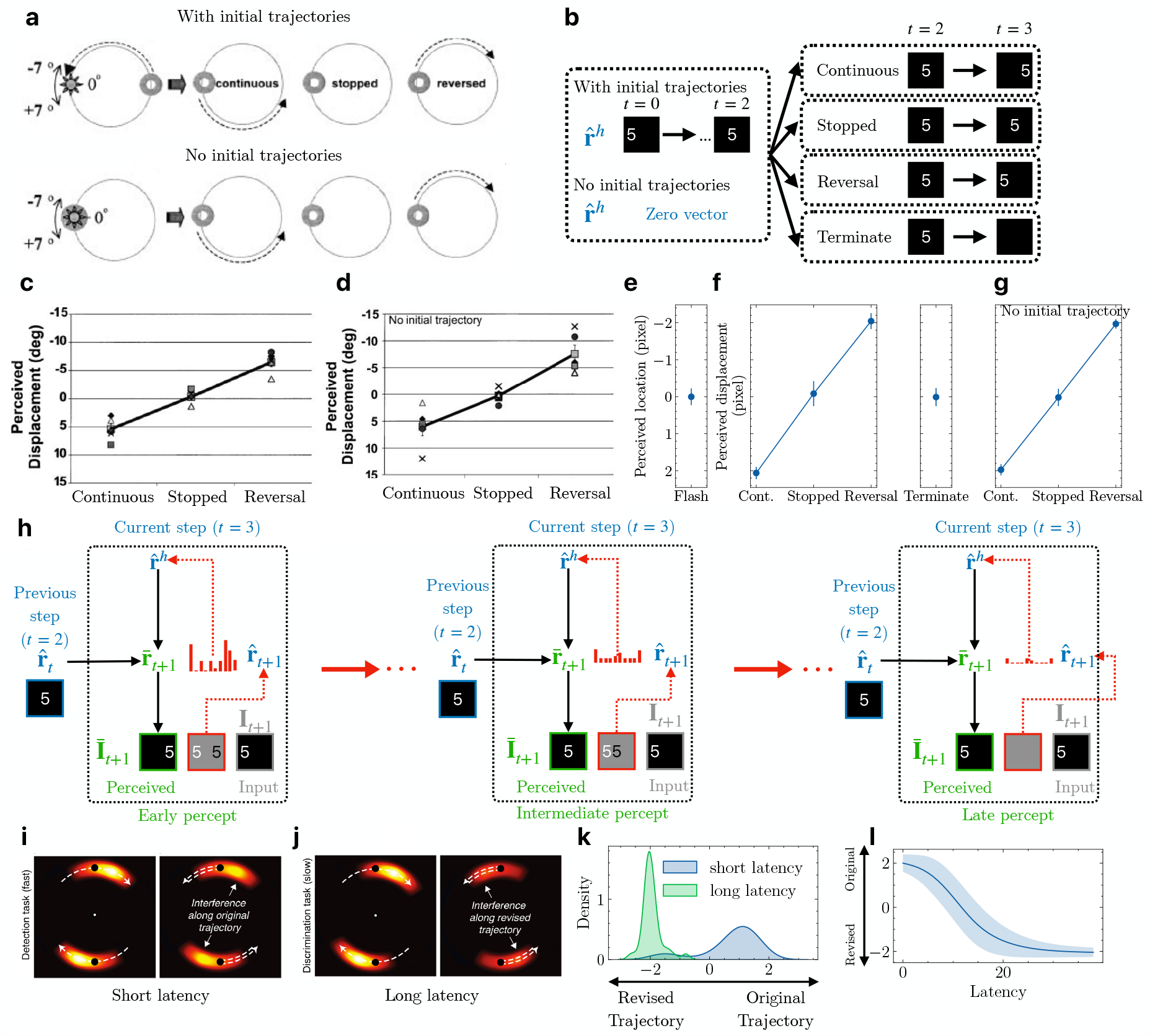
Flash-lag illusion and object representations in apparent motion. **(a)** The flash-lag test conditions used by Eagleman & Sejnowski [26]. The moving ring could have an initial trajectory (top) or no trajectory (bottom). At the time of the flash (bright disk), the ring could move along the initial trajectory, stop, or reverse its trajectory. Adapted from [26]. **(b)** Two test conditions (left) regarding initial trajectories of the moving object (a digit) in the flash-lag experiment with the model, and four test conditions (right) for the moving object. The flashed object was shown at time *t* and turned off at time *t* + 1 (same as the “Terminate” condition). **(c & d)** Psychophysical estimates for human subjects reported by Eagleman & Sejnowski [26] when the moving object had initial trajectories (c) or no initial trajectory (d). **(e)** Perceived location of the flashed object in the DPC model at time *t* + 1. The error bar indicates *±* 1 standard deviation (measured across presentations of different digits). **(f)** Perceived displacement between the moving object (with initial trajectories) and the flashed object in the DPC model for the four test conditions. **(g)** Same as (f) but with no initial trajectory for the moving object. **(h)** Illustration of the prediction-error-driven dynamics of the perception of the moving object in the model when the trajectory reversed at time *t* + 1. Red ellipsis between panels denotes the prediction error minimization process. **(i)** Interference pattern during human apparent motion perception with continuous motion (left) and reversed motion (right) at short latency (fast detection task). Brighter color denotes more interference. Dashed arrows represent object motion direction. Adapted from [29]. **(j)** Same as (i) but at long latency (slow discrimination task) [29]. **(k)** Perceived location of the moving object in the DPC model at time *t* + 1 probed at short versus long latency during prediction error minimization. Positive values denote distance along the original trajectory. Negative values denote distance along the reversed trajectory. Short and long latency correspond to “Early percept” and “Late percept” respectively in part (h). **(l)** Perceived location of the digit at all latencies during the prediction error minimization process in part (h).

We propose that prediction error minimization with a hierarchical temporal representation, as in the DPC model, provides a natural explanation for these predictive and postdictive effects. In a DPC network, the higher-level state **r**^*h*^ predicts entire sequences of lower-level states following the same dynamics (Figure 3). When the dynamics of observations change (e.g., motion reversal), the higher-level state is updated to minimize prediction errors, resulting in a revised state that represents the motion-reversed sequence spanning both past and future inputs. This process corresponds to postdiction in visual processing [28]. For the flash-lag experiment, we predict that the higher-level neurons of a trained DPC network will form a static sequence percept when presented with a flashed object and a directional sequence percept for a moving object, causing perceived lags between the two objects as observed in the flash-lag illusion [30].

We first test these predictions of the DPC model on the experimental conditions used by Eagleman & Sejnowski [26]. In their experiment, the stimuli consisted of a flashed disk and a ring moving in a circle. Before the flash, the ring could have an initial trajectory (Figure 4(a), top) or no initial trajectory (Figure 4(a), bottom). After the flash, the ring could continue moving on its initial trajectory (“continuous”), stop moving (“stopped”), or move on the reversed trajectory (“reversed”). A flash appeared in a seven-degree range that extended above and below the ring on its trajectory. The participants then indicated whether a flashed white disk occurred above or below the center of the moving ring. Positive displacements denoted lags along the initial trajectory of the ring, while negative displacements denoted the reversed direction.

To simulate these testing conditions, we used the Moving MNIST test set and extracted 134 test sequences with consistent leftward or rightward motion. For each of these 134 sequences, we simulated the two test conditions used by Eagleman & Sejnowski [26] (with or without initial trajectory): the higher-level state 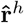 was either inferred from the first three steps (*t* = 0, 1, 2) of the input sequence, or initialized to the zero vector (Figure 4(b) left). For each of these two test conditions, we simulated the four test cases used in [26] regarding the motion of the moving object at the time of the flash (Figure 4(b) right). Note that flashed stimuli correspond to the “no initial trajectory, terminate” condition. We computed the location of a digit as the center of mass of pixel values in the 2D image; the perceived location at time *t* was defined similarly based on the predicted image at time *t*. As Figure 4(e) shows, the perceived location of a flashed object at *t* = 3 strongly overlapped with the physical flashed location at *t* = 2, showing that the prediction errors drove the higher-level state estimates to predict no change in object location for the flashed object. Figure 4(f) shows the perceived displacement between the moving object (with initial trajectories) and the flashed object, computed as the difference in perceived locations at *t* = 3 between the moving object and the flashed object. Positive displacements followed the original trajectory direction and negative displacements followed the reversed direction. The perceived displacements in the model were significantly different in the three test conditions (Figure 4(f) left panel, *p* ≪ 0.001, one-way ANOVA test) and were similar to the psychophysical results reported by Eagleman & Sejnowski (Figure 4(c)). Figure 4(g) confirms that the initial trajectories of the moving object had no effects on the model’s flash-lag illusion, consistent with the reported results (Figure 4(d)) [26]. These results validate the explanation provided by the DPC model on the flash-lag effect: for a hierarchical generative model with representations of sequences, a flashed or stopped/terminated moving object leads to inference of a static object sequence (Figure 4(e)), while continuous or reversed motion leads to inference of a moving object sequence, resulting in the perceived lags along the corresponding directions (Figure 4(f)).

One aspect of motion perception the previous results do not illustrate is the interplay between postdiction and prediction. Hogendoorn et al. investigated this effect in an experiment on apparent motion perception. Participants were instructed to report the detection of a visual cue (short latency task) or differentiate between two visual cues (long latency task) during apparent motion. These visual cues could either be along the apparent motion trajectory or the reversed trajectory. The authors found that upon reversing the apparent motion trajectory, predictive effects dominated perception at short latency (detection task, Figure 4(i)), with the most interference (measured in terms of the participants’ reaction times) along the original motion trajectory. At longer latency (differentiation task, Figure 4(j)), most interference was along the reversed trajectories, indicating that postdictive effects dominated perception.

We hypothesize that the prediction error minimization process of DPC could explain this interplay between prediction and postdiction, as illustrated by Figure 4(h) which depicts the gradient-descent-based optimization process of Figure 1(e) (and Equation 18). Early percepts of the model are dominated by the spatiotemporal prediction using the optimal estimates from the previous step (Figure 4(h) left). When a motion reversal occurs, feedforward prediction errors gradually correct the second-level state (Figure 4(h) middle) until convergence (Figure 4(h) right). Therefore, late percepts in the model correspond to error-corrected spatiotemporal predictions. Note that due to the discrete temporal nature of the DPC model (unit time steps), this process is considered to happen “at” one particular time step (*e*.*g* ., early versus late percept “at” *t* = 3 in Figure 4(h)).

To test this hypothesis, we used the same trained DPC network and probed its percept of the moving object at the time of reversal under the “with initial trajectory, reversal” condition (Figure 4(b)). At short latency (10% of steps into prediction error correction, Figure 4(h) early percept), the perceived locations for the moving object in most test sequences were along the original trajectory, as denoted by positive displacements compared to the final step before reversal (*t* = 2) (Figure 4(k) blue)). At longer latency (90%, Figure 4(h) late percept), the moving object’s perceived locations were flipped and along the reversed trajectory (negative displacements; Figure 4(k) green, *p* ≪ 0.001, *t*-test). This is consistent with psychophysical findings [27, 29] that when the motion of the object unexpectedly reversed, prediction effects were observed at short latency ( ≈ 350 ms, Figure 4(i) right panel, bright color denotes locations of interference due to prediction) while postdictive effects were observed at longer latency ( ≈ 620 ms, Figure 4(j) right panel, bright color denotes locations of interference due to postdiction). Figure 4(l) plots the moving object’s perceived location in our model throughout the error correction process: the perceived location varies smoothly from being along the original direction initially to along the reversed direction at greater latencies. These results make a testable prediction: if probed at an intermediate level of latency (between 350 ms and 620 ms), the maximal interference should overlap with the object’s location at the time of reversal (*i*.*e*., at the black dots in Figure 4(i,j)), as suggested by Figure 4(l).

### Cue-triggered recall and episodic memory

A number of experiments in rodents have shown that the primary visual cortex (V1) encodes predictive representations of upcoming stimuli [1–4, 48]. In one of the first such studies, Xu et al. [1] demonstrated that after exposing rats repeatedly to the same moving dot visual sequence (Figure 5(a)), displaying only the starting dot stimulus triggered sequential firing in V1 neurons in the same order as when displaying the complete sequence (Figure 5(a)). Similar effects have been reported in monkey [35] and human [32–34, 49] visual cortical areas as well.

**Figure 5:**
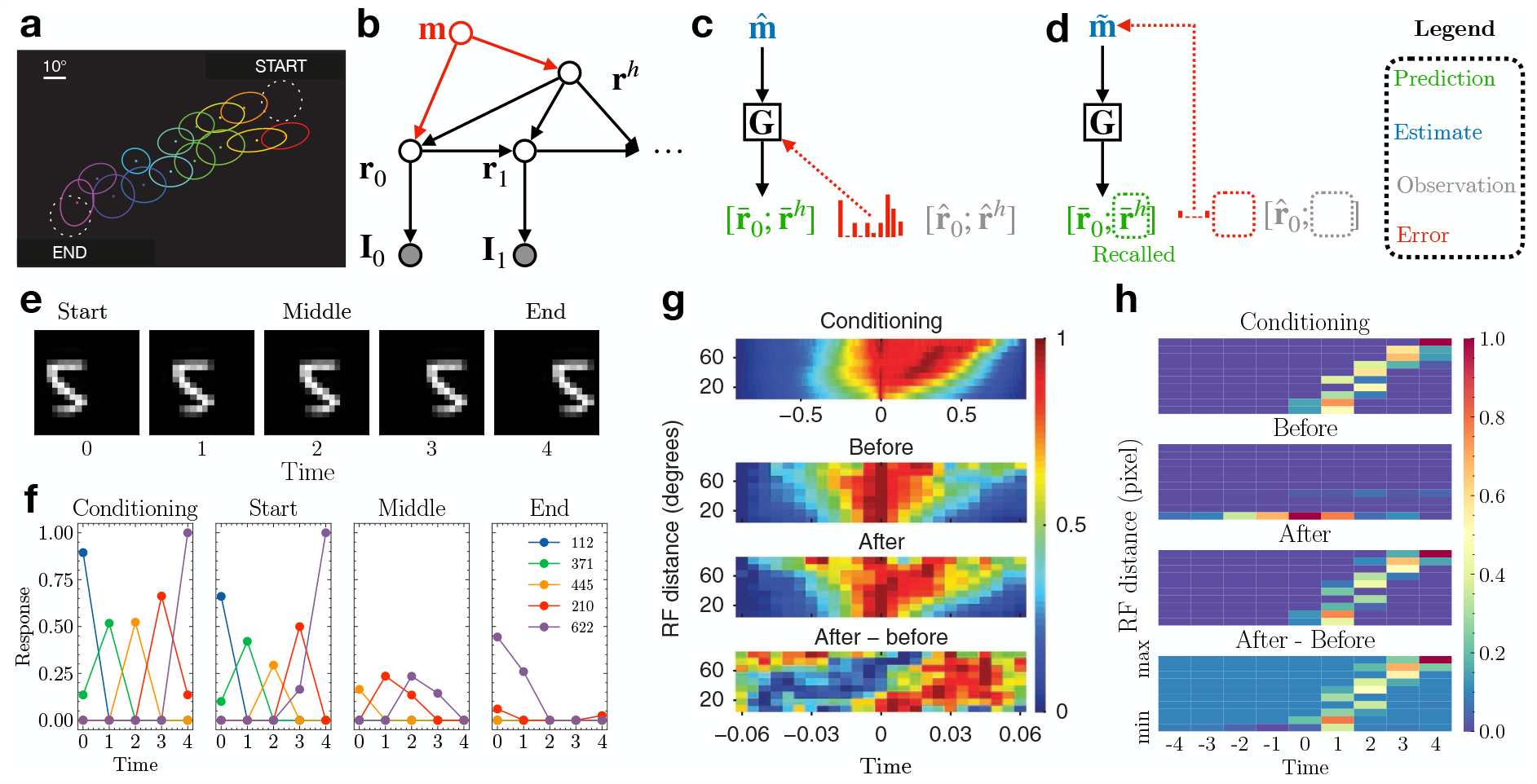
Cue-triggered activity recall in the DPC model. **(a)** The experimental setup of Xu et al. (adapted from [1]). A bright dot stimulus moved from START to END repeatedly during conditioning. Activities of neurons whose receptive fields (colored ellipses) were along the dot’s trajectory were recorded. **(b)** Generative model combining an associative memory and DPC. The red part denotes the augmented memory component that binds the initial content vector **r**_0_ and the dynamics vector **r**^*h*^ to encode an episodic memory. **(c)** Depiction of the memory encoding process. The presynaptic memory activity and postsynaptic prediction error jointly shape the memory weights **G. (d)** Depiction of the recall process. Prediction error on the partial observation 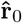 drives the convergence of the memory estimates 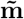 and recalls the higher-level dynamics vector 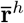 as a top-down prediction. The red dotted box depicts the prediction error between the missing observations for **r**^*h*^ and the prediction 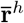; this error is ignored during recall, implementing a form of robust predictive coding [47]. **(e)** The image sequence used to simulate conditioning and testing for our memory-augmented DPC network. **(f)** Responses of the lower-level neurons of the network. Colored lines represent the five most active lower-level neurons at each step. Left to right: neural responses during conditioning, testing the network with a single start frame, middle frame, and end frame. **(g, h)** Normalized pairwise cross correlation of **(g)** primary visual cortex neurons (adapted from [1]) and **(h)** the lower-level model neurons. Top: during conditioning; middle two: testing with the starting stimulus, before and after conditioning; bottom: the differences between cross correlations, “After” minus “Before” conditioning.

The generative model of DPC provides a highly efficient computational basis for episodic memories and sequence prediction. DPC assumes sequences are generated by a factorized representation: a single (lower-level) representation of the content (“what”) provided at the first step and a single (higher-level) representation of dynamics (motion or “where”). These two representations are inferred during sequence perception as the explanations (or causes) of a given input sequence.

It is known that factored information from the visual cortex makes its way, via the medial and lateral entorhinal cortices, to the hippocampus [50]. The hippocampus has been implicated both in the formation of episodic memories [51–53] and in mediating activity recall in the neocortex [54–57] through its outputs to the entorhinal cortex, which in turn conveys this information to downstream areas via feedback connections. Because the DPC model encodes an entire sequence in terms of a single dynamics vector **r**^*h*^ (along with the content **r**_0_ at the first step), it suggests a simple mechanism for storing sequential experiences as episodic memories, namely, storing the vector **r**^*h*^ (along with **r**_0_) in an associative memory, mimicking the role of the hippocampus.

To test this hypothesis, we augmented the DPC model with an associative memory that uses a vector **m** to bind the content vector **r**_0_ with the dynamics of the sequence **r**^*h*^, thereby encoding an episodic memory: the new generative model is shown in Figure 5(b). Given the initial cue **r**_0_ (inferred from the first image frame in the sequence), the associative memory (emulating the hippocampus) recalls the episodic sequence dynamics **r**^*h*^ which modulates the transition dynamics in the DPC network (representing the visual cortex) to complete the sequence recall. Specifically, we added to the trained DPC network another higher level of predictive coding to implement associative memory [10, 58]: the memory vector estimate 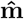 predicts both the content vector 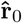 and motion dynamics vector 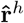 and uses the prediction error to correct itself (Figure 5(c)). Upon convergence of 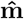, the associative memory network stores this vector by updating its weights **G** using Hebbian plasticity based on presynaptic activity 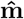 and postsynaptic prediction error [10] (Figure 5(c), see Methods). During recall, the memory estimate 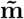 is driven by the 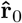 inferred from the cue and the prediction error (Figure 5(d), dashed boxes denote the missing input). The dynamics vector **r**^*h*^ is then recalled as the top-down prediction after 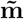 has converged (Figure 5(d), green dashed box). Note that during conditioning, no learning occurs in the DPC network – only the weights of the memory network **G** are optimized to store the episodic memory 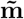 .

We simulated the experiment of Xu et al. [1] using a moving MNIST sequence from the test set shown in Figure 5(e). After conditioning (5 repetitions of the sequence), the network was tested with the starting frame only, the middle frame only, and the end frame only. The lower-level responses 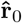 of the DPC network were used to recall the dynamics component 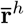 from the memory. The recalled dynamics were then used to predict a sequence of lower-level responses in the DPC network. We found that the lower-level model neurons exhibited cue-triggered activity recall given only the start frame of the sequence (Figure 5(f) Start). Cueing the network with the middle frame triggered weak recall, consistent with findings by Xu et al. (see Figure 3c in Ref. [1]). The end frame did not trigger recall [1]. We found that the sequence recall is cue-specific – when trained with sequences that have distinct digits and dynamics, the DPC network successfully recalled the correct sequence when cued with different starting digits (Figure S2).

Lastly, following the analysis done by Xu et al. [1], we plotted the pairwise cross-correlation of the lower-level model neurons as a function of their spatial RF distances when tested with the starting frame of the sequence (see Methods). As Figure 5(h) shows, the peaks of the correlation showed a clear rightward slant after conditioning, consistent with the experimental results (Figure 5(g)). This indicates a strong sequential firing order in the lower-level model neurons elicited by the starting cue, where neurons farther apart have longer lags in response cross-correlations, a phenomenon that was nonexistent before conditioning (Figure 5(g)). These simulation results support our hypothesis that cue-triggered recall could be the result of the hippocampus, acting as an associative memory, binding factorized sequence representations of content and dynamics from the neocortex and recalling the corresponding dynamics component given the content cue.

### Estimating higher-order transition dynamics with a three-level model

Our results thus far involved a two-level DPC model whose second-level states predicted the first-level state transitions. Since the second-level state is assumed to characterize the *entire* sequence (Figure 1), it cannot predict higher-order transitions such as digits bouncing at the boundaries in the Moving MNIST dataset, which requires different second-level state representations (Figure 3). Here we show that adding a third level allows the DPC model to learn and infer the *transition dynamics of second-level states*, thus capturing a temporally more abstract representation of sequences at the third level.

Figure 6(a) shows the generative model for the three-level DPC model. Just as the second-level states modulate the transition function of the first-level states in the two-level model (Figure 1), the third-level states modulate the transitions of the second-level states. During inference (Figure 6(b)), when the first-level prediction error is larger than a threshold (see Methods), the second-level state transitions to the next state, following the transition function predicted by the current third-level state. The second-level prediction error is conveyed to the third level to correct its state estimate, in the same way as the first-level prediction error corrects the second-level state.

**Figure 6:**
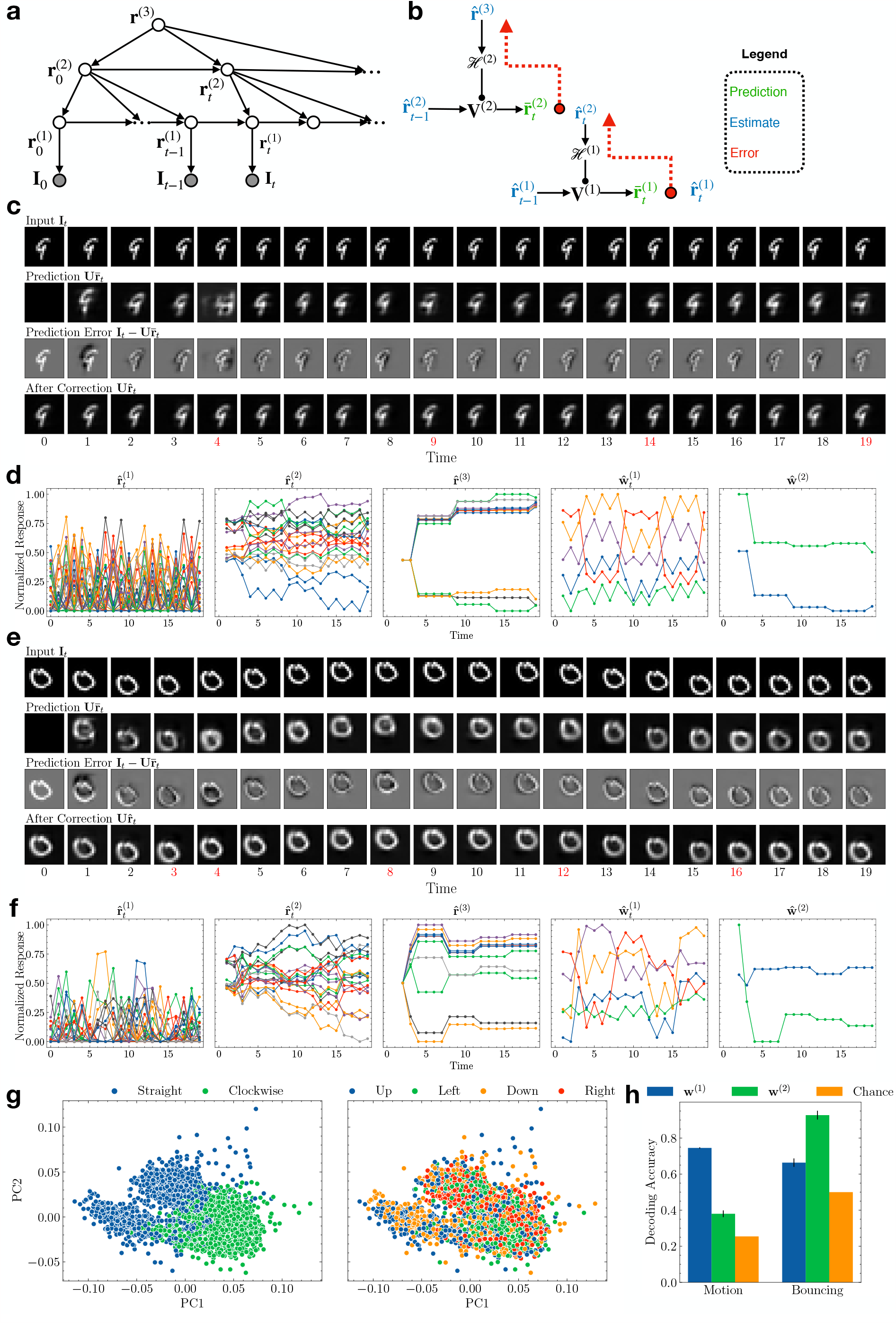
(Previous page.) Three-level DPC model learns progressively more abstract temporal representations. (a) Generative model for three-level DPC. **(b)** Schematic depiction of an inference process. Observation nodes are omitted for clarity. **(c)** Inference for an example Moving MNIST sequence with “straight bouncing” dynamics. Red time steps mark the moments when the first-level prediction error exceeded the threshold, causing the network to transition to a new second-level state (see Methods). For these time steps, the predictions (second row) are by the second-level neurons, while the rest are by the first-level neurons as in Figure 3. **(d)** The network’s responses to the Moving MNIST sequence in (c). Left to right: first-level responses, second-level responses, third-level responses, first-level modulation weights, second-level modulation weights. **(e)** Same as (d) but with “clockwise bouncing” dynamics. **(f)** Same as (d) but for the sequence in (e). **(g)** Third-level responses to the Moving MNIST sequences visualized in the 2D space of the first two principal components. Left: responses colored according to bouncing type; right: responses colored according to motion direction. **(f)** Comparison of decoding performance for bouncing type versus motion direction using the modulation weights generated by the second and third level. Error bars: *±* 1 standard deviation from 10-fold cross validation. Orange: chance accuracies.

The Moving MNIST dataset we used for Figure 3 exhibited only one type of transition dynamics of the second-level states, namely, transitioning from moving left to moving right, or moving up to moving down, and vice versa (henceforth referred to as “straight bouncing” dynamics (Figure 6(c))). To demonstrate that the third-level states can learn different second-level dynamics, we added to the dataset digit sequences with “clockwise bouncing” dynamics to the dataset (for example, a digit moving to the left and hitting the boundary will move upward instead of rightward, and so on (Figure 6(e))). This makes the second-level state transition function ambiguous until the first bouncing event. If the third-level representations learned by DPC capture the second-level transition dynamics, we expect the first large prediction error at the second level (occurring at a boundary) to update the third-level state estimate to represent either straight bouncing dynamics or clockwise bouncing dynamics. Thereafter, the third-level state estimate should remain stable as long as the bouncing type remains the same.

In the following, we use the superscript (*i*) to denote the level *i*. We trained a three-level neural network on the augmented Moving MNIST dataset (with the two types of bounding dynamics discussed above). The network uses two second-level transition matrices 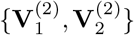 and a top-down network *ℋ*^(2)^ (from the third to the second level), in addition to all the parameters in the two-level model. The first and second-level transition matrices were pretrained (see Methods). Figure 6(c) and (d) show an inference example of the three-layer network on an input sequence with straight bouncing dynamics. Red time steps denote the moments when the first-level prediction errors were larger than the set threshold, causing the second-level neurons to change their activities to transition to the next state (this can be seen as a neural implementation of terminal states in a hierarchical HMM [59] (see Methods)). As seen in the second row in Figure 6(c), at the first bouncing event (*t* = 4), the second-level prediction was not accurate; the third-level neural responses were updated to minimize this prediction error (see Figure 6(d)). For the rest of the sequence, the predictions are accurate at the bouncing events (*t* = 9, 14, 19) and the third-level neural responses remained stable. The panels in Figure 6(d) show an increase in response stability and timescale from the first to the third-level neural responses (first three panels), as well as in the modulation weights that define the first and second-level transition dynamics (last two panels). Figure 3(e) and (f) show a different example with clockwise bouncing dynamics. Similar to the example above, the third-level responses showed notable changes at times *t* = 3 and 4 but remained stable for the rest of the sequence. Comparing the second-level modulation weights in Figure 6(d) and (f), it is clear that the third-level DPC neurons estimated different bouncing types and generated opposite modulation strengths for the two types of sequences.

We performed PCA on the third-level responses **r**^(3)^ obtained at the end of the Moving MNIST sequences (*t* = 19) in the test set. Figure 6(g) visualizes these responses in the space of their first two principal components (PCs), colored either by bouncing dynamics type (left) or moving direction (right). The third-level responses form clusters according to the bouncing dynamics type but not motion direction. We then used SVMs to decode the bouncing dynamics type or motion direction from **w**^(1)^ and **w**^(2)^, the weights predicted by **r**^(2)^ and **r**^(3)^ respectively. As shown in Figure 6(h), using **w**^(2)^, the classifier yielded 92.7% 10-fold cross-validated classification accuracy on the two bouncing types (chance accuracy: 50.0%). Using **w**^(1)^ resulted in significantly less classification accuracy for bouncing type (66.4%, *p* ≪ 0.001, *t*-test). In contrast, decoding accuracy for the four motion directions (chance accurary: 25.4%) was significantly higher using **w**^(1)^ (74.5%) compared to using **w**^(2)^ (38.0%, *p* ≪ 0.001, *t*-test). These results show that the three-level DPC model succeeded in learning a temporal hierarchy, with the third-level states encoding the longest timescale feature, *i*.*e*. the type of bouncing dynamics, by modulating the transition function of the second-level states, which in turn encoded intermediate timescale features (motion direction).

## 3 Discussion

Our results show that dynamic predictive coding (DPC) can learn hierarchical temporal representations of sequences through top-down modulation of lower-level dynamics. Specifically, we showed that by minimizing prediction errors on image sequences, a two-level DPC neural network develops V1-like separable and inseparable space-time receptive fields at the lower level [25], and representations encoding sequences at a longer timescale at the higher level [6, 8]. The trained DPC network provides a normative explanation for the flash-lag effect [26] and accounts for both prediction and postdiction in visual motion processing [27, 28, 60]. The temporal abstraction of sequences in a DPC network suggests a new mechanism for storing and retrieving episodic memories by linking the DPC network to an associative memory, emulating the interaction between the neocortex and the hippocampus. We show that such a memory-augmented DPC model explains cue-triggered activity recall in the visual cortex [1]. Finally, we show that the top level of a three-level DPC network captures the higher-order temporal statistics encoding the transition dynamics of the second-level states, which in turn capture the temporal statistics of the first-level states. Taken together, the hierarchical temporal representations learned by DPC, ranging from the lowest-level space-time representations similar to those observed in visual cortical simple cells (Figure 2), through the intermediate-level representations of steady motion (Figure 3), to the highest-level representation of how such motion changes over a longer timescale (Figure 6), emulate the spatiotemporal representations observed in visual cortical hierarchies, particularly along the dorsal visual pathway [61].

The key to the DPC model’s ability to capture lower-level dynamics with relatively stable higher-level response vectors **r**^*h*^ is the top-down modulation of transition dynamics of entire lower-level state sequences, using the weights **w** generated by the higher level. There has been increasing interest in neuroscience in the role of modulatory inputs (*e*.*g*., encoding top-down contextual information) in shaping the dynamics of recurrent neural networks in the brain [62–64]. The DPC model ascribes an important role to these modulatory inputs in enabling cortical circuits to learn temporal hierarchies. The neural implementation used in this paper can be seen as top-down feedback (**w**) targeting the distal apical dendrites of lower-level pyramidal neurons, thereby changing their gain (Figure 1(c)). Such a mechanism, which has been shown to be possible experimentally [16, 17, 39, 65], can also modulate perceptual detection [66]. Note that although we chose to model the top-down influence as multiplicative gain modulation, it would be theoretically equivalent to model it as an additive component or concatenate it as an extra input for prediction (*e*.*g* ., predicting first-level transitions as 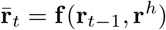, where **f** is a multi-layer perception). However, such an implementation may be less efficient (in terms of the number of parameters required to reach the same level of performance) under certain conditions, compared to a hypernetwork-based implementation [67] such as our implementation based on multiplicative gain modulation.

Some of the first models of spatiotemporal predictive coding focused on signal processing in the retina and LGN [68, 69]. Other models for sequence processing, such as sparse coding [20, 38] and independent component analysis [18], have been shown to produce oriented space-time receptive fields from natural image sequences, but these models require the entire image sequence to be presented as a single vector input, which is hard to justify biologically; they also do not explicitly model the temporal dynamics between images and therefore, cannot make predictions into the future given a single input. A previous spatiotemporal predictive coding model based on Kalman filtering [40] did incorporate state transitions but the model was not hierarchical and was not shown to generate cortical space-time receptive fields. Our model bears some similarities to slow feature analysis which extracts slowly varying features from sequences of stimuli but it does not learn the transition dynamics between time steps [19, 21, 22]. DPC on the other hand learns a generative model that generates entire sequences, with the assumption that the transition dynamics do not change within a sequence (a “slow” feature). Object identity remains in the lower-level representations of DPC (Figure 3(f)). From a learning perspective, Luczak et al. [70] propose that single neurons predicting their future activity at a fixed delay could also serve as an effective learning mechanism.

Recent advances in deep learning [71] have spurred several efforts to learn spatiotemporal hierarchies from sensory data. Lotter et al. developed a deep learning model called “PredNet” for learning a hierarchical predictive coding-inspired model for natural videos [72, 73]. After training, the model was shown to produce a wide range of visual cortical properties and motion illusions. However, in PredNet, higher-level neurons predict lower-level prediction *errors* rather than neural activities or dynamics, making it unclear what the underlying generative model is. It is also unclear if PredNet learns a temporal response hierarchy as found in the cortex. A different model, proposed by Singer et al. [23] and later extended to hierarchies [24], is trained by making higher layers predict lower layer activities: after training, model neurons in different layers displayed different levels of tuning properties and direction selectivity similar to neurons in the dorsal visual pathway. However, similar to the sparse coding and ICA models discussed above for spatiotemporal sequences, the Singer et al. model also requires a sequence of images to be presented as a single input to the network, and the hierarchy of timescales is hard-coded (higher-level neurons predict future lower-level neural activities by receiving a fixed-length chunk of neural activities as input). The above models also do not provide explanations for postdiction or episodic memory and recall.

Many experimental studies have shown an increase in temporal representation stability and response timescales as one goes from lower-order to higher-order areas in the visual and other parts of the cortex [6–9, 74, 75]. Most computational models have studied this phenomenon through mechanistic rate-based models with parameters based on connectivity data [76, 77] or spiking network models [78]. Kiebel et al. [79] proposed a model where second-level states generate a single parameter for the first-level Lorenz attractor as the slower “sensory cause” parameter. DPC generalizes this model by assuming higher-level states fully determine the lower-level transition function by predicting the transition dynamics of lower-level states. Under this formulation, temporal hierarchies emerge naturally as a consequence of the neocortex learning from temporally structured data (*e*.*g*., stable dynamics in short time windows). This view is consistent with findings that response timescales are functionally dynamic and could expand for cognitive tasks such as working memory [80].

Previous normative models of postdiction in visual processing often relate the effect to the concept of Bayesian smoothing (or backward message passing) [26, 81]. We have shown that a trained two-level DPC network with higher-level sequence representations also exhibits postdictive effects without the need for smoothing. In the event of a temporal irregularity (*e*.*g*., an unexpected motion reversal), the higher-level state in the DPC network is updated to reflect a new revised input sequence, naturally implementing postdiction through online hierarchical Bayesian filtering (Figures 1 and 3). Our flash-lag simulation results are consistent with the Bayesian filtering model from Khoei et al. [82] showing that the flash-lag effect can be produced through an internal model that explicitly represents object velocity. The higher-level sequence representation in the DPC model supports an implicit (and more generalized) representation of velocity and reproduces the same internal dynamics of the “speed” estimate at motion reversal (compare Figure 4(h) with Figure 6 in [82]). It is worth noting that the trained DPC network learned to predict no motion (static sequence) for the flashed object even though it was never trained on static object sequences and did not assume a prior of zero speed [82]. This emergent property was also seen in PredNet, which learned to predict relatively little motion for a flashed bar stimulus [73].

The higher-level sequence representations of DPC, when combined with an associative memory, support the formation of episodic memories and cue-triggered activity recall [1, 3, 32–35]. The associative memory in our model forms an episodic memory by binding the inferred content representation and dynamics representation from the DPC network during conditioning. When an initial portion of the sequence is presented during testing, the stored episodic memory is retrieved, generating the dynamics component which modulates the lower-level network to enable full recall of the sequence. Though previously considered to only require V1 plasticity [1, 3], sequence learning is severely impaired in mice with hippocampal damage [83]. Coordinated activity between V1 and the hippocampus has also been found in human V1 during recall [49]. These experimental results support the involvement of the hippocampus in sequence learning, consistent with our model. Overall, the memory-augmented DPC model offers a highly efficient computational basis for forming and recalling episodic memories [57, 84], where a single representation of content and transition dynamics from all sensory areas of the neocortex can be bound together as a memory and later retrieved upon receiving partial input.

In our three-level DPC model, the second-level state (under the influence of the current third-level state) predicts the next second-level state only when the first-level prediction errors are larger than an estimated threshold. This can be seen as a neural (and continuous-valued) implementation of “terminal states” in hierarchical hidden Markov models (HHMMs) [59]. In an HHMM, when a terminal state is reached at a lower level, the corresponding sub-HMM is deemed to be completed and the higher-level state then transitions to the next higher-level state (which activates the next sub-HMM at the lower level). In our hierarchical DPC model, small first-level prediction errors are resolved locally between the first and second level, indicating a continuing sub-sequence. When the error exceeds the threshold, the sub-sequence ends and the second-level transitions are activated. Any second-level prediction errors are resolved between the second and third level through third-level state inference. We used post-hoc estimation of the error threshold after training but future work could attempt to estimate the threshold online in terms of the inverse variance or “precision” of prediction errors [85]). Additionally, second-level transitions could also correlate with the spatial information in the videos (*e*.*g* . bouncing only happens when the digit is near the boundary). Models whose second-level states depend on both the previous first- and previous second-level states could learn this type of transition [86].

The DPC model can be extended to action-conditioned prediction and hierarchical planning (see, e.g., [87] for initial steps in this direction). There is a growing body of evidence that neural activity in the sensory cortex is predictive of the sensory consequences of an animal’s own actions [2, 4, 13, 48, 88]. These results can be understood in the context of a DPC model in which the transition function at each level is a function of both a state and an action at that level, thereby allowing the hierarchical network to predict the consequences of actions at multiple levels of abstraction [87]. Such a model allows probabilistic inference to be used not only for perception but also for hierarchical planning, where actions are selected to minimize the sensory prediction errors with respect to preferred goal states. Such a model is consistent with theories of active inference [11] and planning by inference [89–93], and opens the door to understanding the neural basis of navigation and planning [9, 94, 95] as an emergent property of prediction error minimization.

## 4 Methods

### Hierarchical generative model

We assume that the observation **I**_*t*_ ∈ ℝ^*M*^ at time *t* is generated by a lower-level latent variable **r**_*t*_ ∈ ℝ^*N*^ . The latent variable **r**_*t*_ is generated by the previous step latent variable **r**_*t*−1_ and the higher-level latent variable, 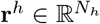. Together, the generative model factorizes as follows:

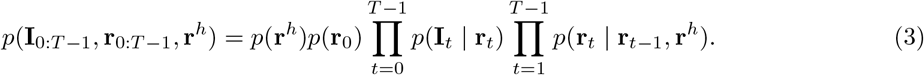

Each component of the factorization is parameterized as follows:

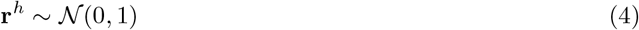

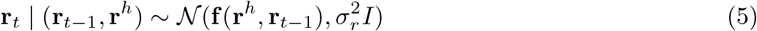

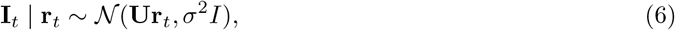

where *𝒩* denotes the normal distribution and *I* denotes the identity matrix. The mean 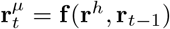 is given by:

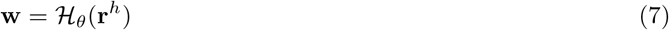

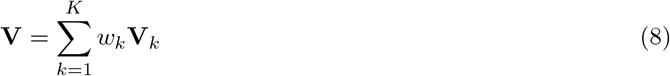

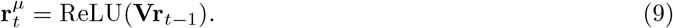

Here, *ℋ*_*θ*_ is a function (neural network) parameterized by *θ*.

To sum up, the trainable parameters of the model include spatial filters **U**, *K* transition matrices **V**_1_, …, **V**_*K*_, and the neural network parameters *θ*. The latent variables are **r**_0:*T* −1_ and **r**^*h*^. See Supplementary Information for a more detailed description of the model architecture.

### Prediction error minimization

Here, we derive the loss function used for inference and learning under the assumed generative model. We focus on finding the *maximum a posteriori* (MAP) estimates of the latent variables using a Bayesian filtering approach. At time *t*, the posterior of **r**_*t*_ conditioned on the input observations up to time *t*, **I**_0:*t*_, and the higher-level variable **r**^*h*^ can be written as follows using Bayes’ theorem:

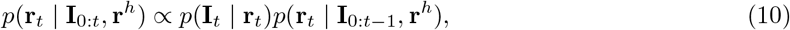

where the first term on the right-hand side is the likelihood function defined by Equation 6. The second term is the posterior of **r**_*t*_ given input up to the previous step and the higher-level state:

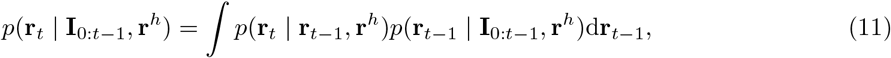

where the first term inside the integral is the lower-level transition dynamics defined by Equation 5. Note that the parameterization of the transition distribution is generated by the higher-level latent variable as specified by Equations 7-9.

Putting Equation 10 and 11 together, we get

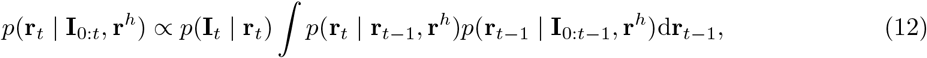

which defines a recursive way to infer the posterior of **r**_*t*_ at time *t*. In this model, we only maintain a single point (MAP) estimate of the posterior at each time step, so we simplify the posterior distribution *p*(**r**_*t*−1_ | **I**_0:*t*−1_, **r**^*h*^) as a Dirac delta function:

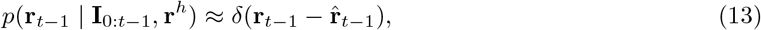

where 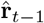 is the MAP estimate from the previous step. Now we can further simplify Equation 12 as

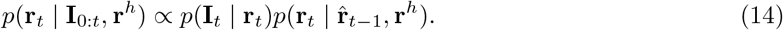

This gives the posterior of all the latent variables at time *t* as

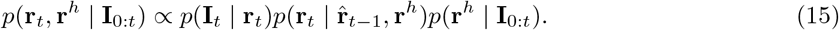

We can find the MAP estimates of the latent variables by minimizing the negative log of Equation 15. Substituting the generative assumptions (Equations 4 to 6), we get:

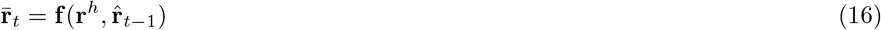

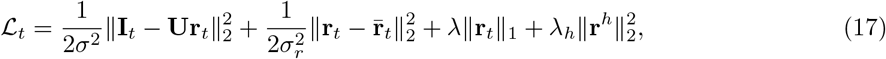

where *λ* and *λ*_*h*_ are the sparsity penalty for **r**_*t*_ and the Gaussian prior penalty for **r**_*h*_, respectively. Note that we approximate *p*(**r**^*h*^ **I**_0:*t*_) with the unconditional prior *p*(**r**^*h*^) so that at each step the dynamics are estimated using only the local pairwise transition and the prior. Using Equation 17, we compute the MAP estimate of **r**_*t*_ at time *t* as

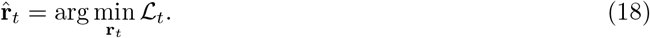

At each time step, we update the current estimate of **r**^*h*^ to minimize ℒ_*t*_ as well:

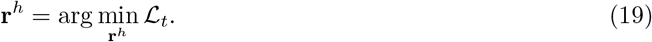

To begin the recursive estimation (without the temporal prediction from the previous step), we compute the MAP estimate of the first step latent variable **r**_0_ using the following reduced loss

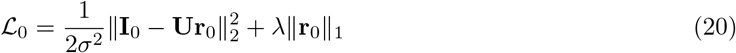

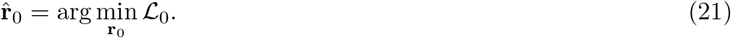

The parameters of the model can be optimized by minimizing the same prediction errors summed across time and averaged across different sequences, using the MAP estimates of the latent variables. See Supplementary Information for detailed pseudocode describing the inference and learning procedure.

### Data and preprocessing

For the natural video dataset, we extracted 65520 image sequences from a YouTube video (link here) recorded by a person walking on a forest trail (image size: 16 *×* 16 pixels, sequence length: 10 frames (≈ 0.35 seconds, uniformly sampled in time). The image sequences do not overlap with each other spatially or temporally. Each sequence was spatially and temporally whitened to simulate retinal and LGN processing following the methods in Olshsausen & Field [38] and Dong & Atick [41]. 58,968 sequences were used to train the model and the remaining 6,552 were reserved for testing.

For the Moving MNIST dataset [45], we used 10000 image sequences (image size: 18 *×* 18 pixels, sequence length: 10 frames), each sequence containing a fixed digit moving in a particular direction. The motion of the digits was restricted to upward, downward, leftward, or rightward directions. When a digit hit the boundary, its motion direction was inverted (leftward to rightward, upward to downward, and vice versa). No whitening procedures were performed on the MNIST sequences. 9,000 sequences were used to train the model and the remaining 1,000 were reserved for testing.

### Reverse correlation for computing space-time receptive fields

The reverse correlation stimuli for deriving the space-time receptive fields (Figure 2) were extracted from the same natural video data but without any spatial and temporal overlapping with the training and test set. We used 50 continuous image sequences with 80,000 steps ( ≈ 47 minutes, spanned the same time range but no spatial overlaps) and computed the space-time receptive fields (STRFs) as the firing-rate-weighted average of input frame sequences of length seven ( ≈ 250 ms), across time and sequences.

Formally, let 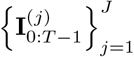 be the *J* stimulus sequences of length *T* and let *τ* be the length of the STRFs (here, *J* = 50, *T* = 80, 000, *τ* = 7). For each neuron *i*, its space-time receptive field STRF_*i*_ has dimensions *M × τ*, where *M* is the dimensionality of a single image frame vector (here, *M* = 100 after vectorizing the 10 *×* 10 image frame). We compute the STRF of neuron *i* as follows

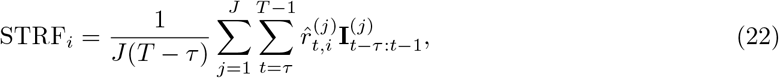

where 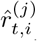 is the predicted firing rate of neuron *i* at time *t* in sequence *j* and 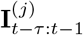 is the image sequence from time *t* − *τ* to *t* − 1 in sequence *j*. This procedure is analogous to the calculation of the spike-triggered average widely used in neurophysiology [43]; in this case, we computed the average of input sequences weighted by the activity 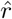 caused by the sequence.

### Autocorrelation for quantifying timescales

Since our model responds deterministically to the same sequence (MAP estimation), we cannot follow the exact approach of Murray et al. [6] that relies on across-trial variability to the same stimulus. We computed the autocorrelation of single neuron responses to natural videos and Gaussian white noise sequences. We averaged the single-neuron autocorrelations across the lower or higher level, and trials. Formally, let *r*_*t,i,j*_ be the response of neuron *i* at time *t* during trial *j*. We computed the autocorrelation with lag *k* as follows:

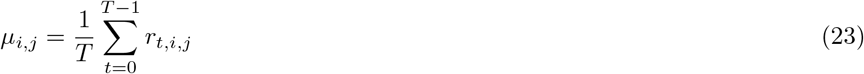

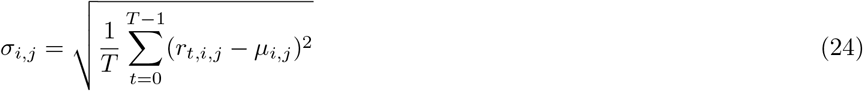

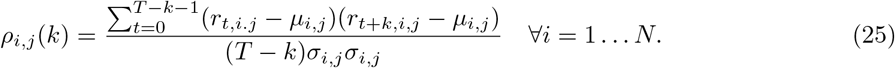

To compute the autocorrelation for an entire population at lag *k*, we took the average of the autocorrelations across all *N* neurons and *J* trials:

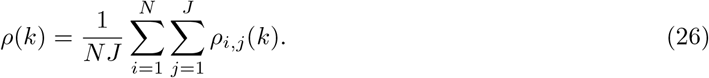

For the results shown in Figure 3(b) and (c), we choose *J* = 500, *T* = 50 and computed the autocorrelation with lag *k* = 0 … 9 for the lower- and higher-level neurons. White noise pixels were i.i.d. samples from (0, 0.0075). We also computed the autocorrelation for both levels with natural videos selected from the same stimuli used for the reverse correlation analysis (note though with natural videos the stationary mean and variance assumption is less valid). To quantify the timescale of the autocorrelation decay, we fitted an exponential decay function *ρ*_fit_(*k*) = *a* exp( − *k/τ* ) + *b* to the autocorrelation data on each level (through Scipy `optimize.curve_fit` function), where *a, b*, and *τ* are fitted parameters of the function and *τ* represents the response timescale following the definition by Murray et al. [6].

### Flash-lag and postdiction simulation

We extracted five-step sequences that have a consistent leftward or rightward motion from the Moving MNIST test set sequences (134 sequences in total, see Figure 5(e) for an example). To simulate the test conditions used by Eagleman & Sejnowski [26], we either used the first three steps of the sequences to infer a motion (dynamics) estimate 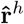 (conditions with initial trajectories), or initialized 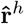 to zero vectors (conditions without initial trajectories). Depending on the test condition, the moving object stimulus at *t* = 3 could move following the original trajectory (“Continuous”), remain at the same location (“Stopped”), move in the reversed trajectory (“Reversed”), or disappear shown an empty frame (“Terminated”), shown in Figure 4(a). The stimuli for simulating flashes correspond to the “no initial trajectory, terminate case”.

The model’s percept of either the moving object or the flashed object at *t* = 3 was computed as the top-down spatiotemporal prediction of the image after correcting the prediction error at *t* = 3:

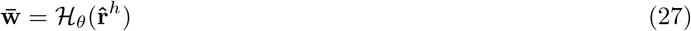

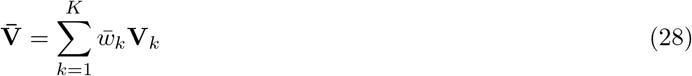

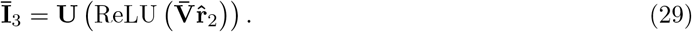

Here, *ℋ*_*θ*_ and **V**_1_, …, **V**_*K*_ are the parameters defined in Equations 7 and 8, 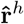 is the optimal higher-level estimate at *t* = 3 (Equation 19), and 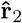 is the optimal lower-level estimate at *t* = 2. We computed the location of the percept as the center of mass of the percept image **Ī**_3_. The displacement in percept between the moving object and the flashed object was calculated as

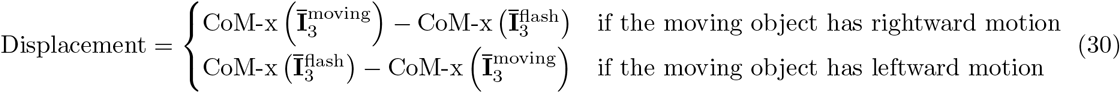

where CoM-x(**I**) returns the horizontal location (*x* dimension) of the center of mass of **I**. Therefore, a positive displacement is along the original trajectory of the moving object, while a negative displacement is along the reversed trajectory.

To compute the plots shown in Figure 4(g)&(h), we used the “with initial trajectory, reversal” test condition. The displacement was computed as in Equation 30 but 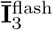 was replaced by the input image **I**_2_ at *t* = 2, at every value of **r**^*h*^ through the error correction process at *t* = 3 (Equations 27 to 30, see Figure 4(b) Perceived versus Input).

### Sequence learning and recall simulation

To simulate the sequence learning experiment of Xu et al. [1], we used a five-step sequence extracted from a Moving MNIST test set sequence (Figure 5(e)). We augmented the hierarchical generative model of DPC with an associative memory layer **m**, which implements predictive coding of the joint higher-level state **r**^*h*^ and the lower-level state **r**_0_ through synapses **G** [10, 58] (see Supplementary Information for model details):

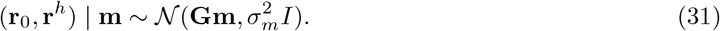

The memory layer was trained separately (the DPC network weights were fixed during conditioning and recall) by minimizing the prediction error:

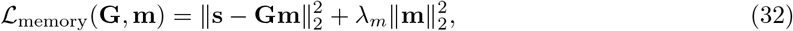

where 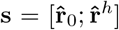 is the concatenated lower- and higher-level state estimates from DPC and *λ*_*m*_ is the regularization penalty on **m**.

During memory encoding, 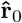 and 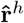 were estimated by the two-level DPC network from the sequence shown in Figure 5(e). Then the memory vector **m** was estimated by gradient descent on Equation 32, yielding the optimal estimate 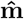 :

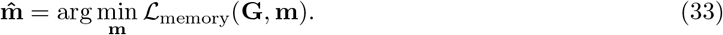

The remaining prediction error drives rapid synaptic plasticity in **G** through gradient descent on the same equation (Figure 5(c)):

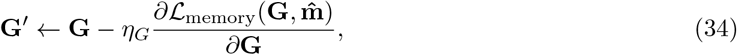

where *η*_*G*_ is the learning rate for **G**. During conditioning, we updated the synaptic weights five times using Equation 34. During recall, given the cue for the beginning of the sequence, the full memory vector **m** is retrieved by minimizing the prediction error with respect to the initial lower-level state **r**_0_ portion of the stored memory only [58] (Figure 5(d)):

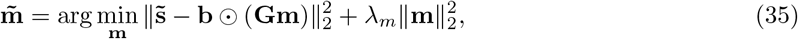

Where 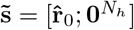 denotes the partial input (visual cue representing the first element of the sequence), ⊙ denotes element-wise multiplication, and 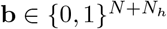 is a binary mask:

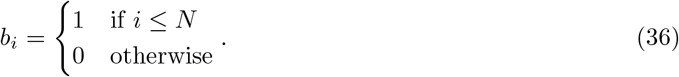

The rest of the sequence was retrieved by retrieving the stored higher-level state 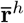 (the dynamics of the sequence) as the top-down prediction through 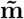 (Figure 5(d)). Once the dynamics were retrieved, we tested sequential recall in the network by predicting an entire five-step sequence using the lower-level vector 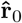 from the visual cue and the retrieved dynamics vector 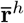 from the memory (Equations 7 to 9). We tested recall in the network using three different visual cues: the starting frame (*t* = 0), the middle frame (*t* = 2), or the end frame (*t* = 4) (see Figure 5(e)&(f)). The cross correlation plot was computed following the same procedure as the one described in Xu et al. [1] (Figure 5(g)&(h)).

### Three-level DPC model

To make the level notation clear, we denote the first and second-level states **r** and **r**^*h*^ from the two-level model as **r**^(1)^ and **r**^(2)^ in the three-level model, and denote the highest (third-level) state as **r**^(3)^. We use superscripts to denote level and subscripts to denote time, unless noted otherwise. The trainable parameters for the three-level model include those for the two-level model as well as two second-level transition matrices 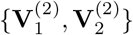 and the third-level top-down network 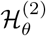 that generates the second-level modulation weights.

### Pretraining the second-level transition matrices

Before training the three-level network, we first pretrained two two-level DPC networks, each with a single transition matrix **V**^(2)^ on Moving MNIST sequences with either straight bouncing type or clockwise bouncing type. We performed inference on the second-level states with the following loss function:

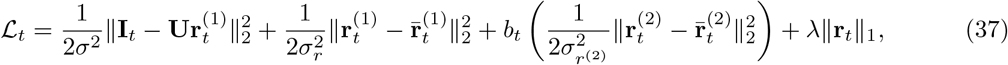

where 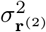 is the variance of second-level prediction errors, *b*_*t*_ ∈ *{*0, 1*}* is a binary mask that equals 1 if the first-level prediction error is larger than a threshold estimated from the training set and 0 otherwise (Figure S3), and 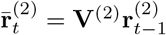 is the predicted second-level state. We learned **V**^(2)^ by gradient descent on the same loss as in Equation 37, summed across time and averaged across sequences, using the MAP estimates of the latent variables.

### Three-level model training

We inferred the second- and third-level states using the same loss function (Equation 37), with the predicted second-level state 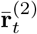 defined as

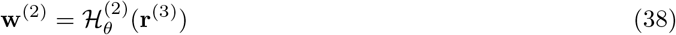

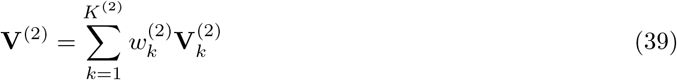

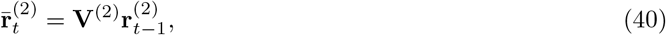

where *K*^(2)^ = 2 is the number of second-level transition matrices. Comparing this definition with Equations 7 to 9, it is easy to see that the third-level top-down prediction is recursively defined in the same way as the top-down prediction in the two-level model. After obtaining the MAP estimates of the second- and third-level states, we learned the third-level top-down network 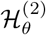 by gradient descent on the same loss. See Supplementary Information for detailed pseudocode describing the inference and learning procedure for the three-level model.

## Data Availability

Data and code for reproducing all simulations in the paper are available at https://github.com/lpjiang97/dynamic-predictive-coding

## Acknowledgements

The authors would like to thank Ares Fisher, Dimitrios Gklezakos, Prashant Rangarajan and Vishwas Sathish for discussions related to hypernetworks and predictive coding. LPJ thanks Daogao Liu for inspiring discussions on the modeling aspects of the paper. This research was supported by National Institutes of Health Grant no. 1UF1NS126485-01, National Science Foundation (NSF) EFRI Grant no. 2223495, Defense Advanced Research Projects Agency (DARPA) Contract no. HR001120C0021, a UW + Amazon Science Hub grant, a Weill Neurohub Investigator grant, a Frameworks grant from the Templeton World Charity Foundation, and a Cherng Jia and Elizabeth Yun Hwang Professorship to RPNR. The opinions expressed in this publication are those of the authors and do not necessarily reflect the views of the funders.

## Supplementary Information

### Hypernetworks and neural gain modulation

A hypernetwork ℋ_*ϕ*_ is a neural network parameterized by synaptic weights and bias parameters *ϕ* that generates the parameters (weights and bias) of another “primary” neural network **f** [15]:

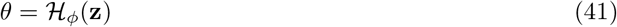

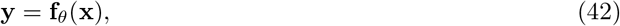

where **z** and **x** are the inputs to the hypernetwork and the primary network, respectively, and **y** is the output of the primary network. Therefore, the responses of the primary network to the same input **x** can be different, depending on the parameters *θ* generated by the hypernet. The input **z** to the hypernet can be a top-down input, a task input or context, an external input, or even past inputs **x**. The hypernet model thus provides a general framework for one neural network (hypernet) modulating the function being computed by another neural network (primary net).

A special case of the hypernet model corresponds to neural gain modulation – a phenomenon where external or internal drives modulate the input-output (I/O) relationship of a neuron [16, 17]. Typically, gain modulation is implemented through multiplicative (altering the slope of the I/O relationship) or additive (shifting the I/O relationship) mechanisms on either the input or the response [16, 39, 96]. These multiplicative and additive components can be seen as the output of the hypernetwork fed to a primary network with fixed parameters *θ*. For example, the recurrent network model proposed by Stroud et al. (Equations 1 in [63]) can be written as (following our notation)

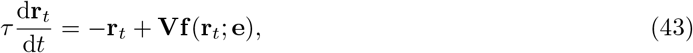

where *τ* is the time constant, **r**_*t*_ is the neural activity vector at time *t*, **V** is the recurrent weight matrix, and **e** is the gain modulation that works as a pointwise multiplicative factor on the input (*f*_*i*_(*r*_*i*_; *e*_*i*_) ≈ tanh(*e*_*i*_*r*_*i*_), see Equation 2 in Ref. [63]). Here, **e** can be seen as the output of a hypernetwork *ℋ*_*ϕ*_ that modulates the recurrent network with weights *θ* = **V**.

Similarly, the DPC generative model for hierarchical temporal prediction through top-down modulation of recurrent connections (Equations 7 to 9) can be implemented using the recurrent network model:

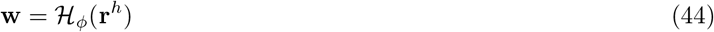

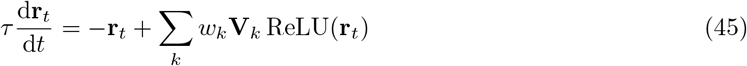

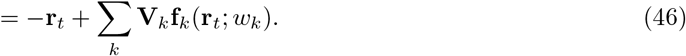

where **f**_*k*_(**r**_*t*_; *w*_*k*_) = *w*_*k*_ ReLU(**r**_*t*_) is the input modulation function which modulates the inputs ReLU(**r**_*t*_) to group *k* of neurons multiplicatively with the weight *w*_*k*_. Therefore, comparing Equation 43 with Equation 46, the gain modulation model of Stroud et al. [63] uses *per-neuron* multiplicative gain modulation while DPC generates a *more diffuse* top-down multiplicative gain modulation targeting *groups of lower-level neurons*, with the same top-down modulation strength *w*_*k*_ for group *k*. Such a model for top-down modulation is consistent with previous work suggesting that cortical feedback connections are more diffuse than feedforward connections [97–99] (but see also [100]).

### Summary of the DPC generative model

Table S1 summarizes the parameters and values used in the DPC generative model. The top-down dynamics prediction network ℋ (a hypernetwork) is a multi-layer perceptron (MLP) as follows. The first layer of the MLP contains 10 hidden neurons, followed by a LayerNorm layer [101] and an ELU activation function [102]. The last two layers of the MLP are comprised of (1) a linear layer with 10 hidden neurons, and (2) an output layer that maps this hidden activity to **w** ∈ ℝ^*K*^ .

### Inference and learning

Table S2 summarizes the optimizers and learning rates used for inference and learning in DPC. Algorithm S1 describes the inference (finding the MAP estimates of the latent variables) and learning (parameter estimation) process.

### Training results

Figure S1(a) visualizes the evolution of test set loss as a function of training epochs. Besides results using five transition matrices (*K* = 5), we additionally trained models with *K* = 1, …, 6, each with eight random initialization of weights. DPC with *K* = 1 is equivalent to a one-level Kalman filter model by Rao [40] with an additional sparseness constraint on neural activities [38]. As shown in Figure S1, all trained models converged after around 50 training epochs. DPC models with two-level architectures (*K >* 1) performed significantly better on the test sets. Such improvement in test set performance saturated as *K* increased. Figure S1(b) shows test set loss as a function of *K* for natural videos and the moving MNIST dataset.

#### Algorithm S1 Inference & learning process

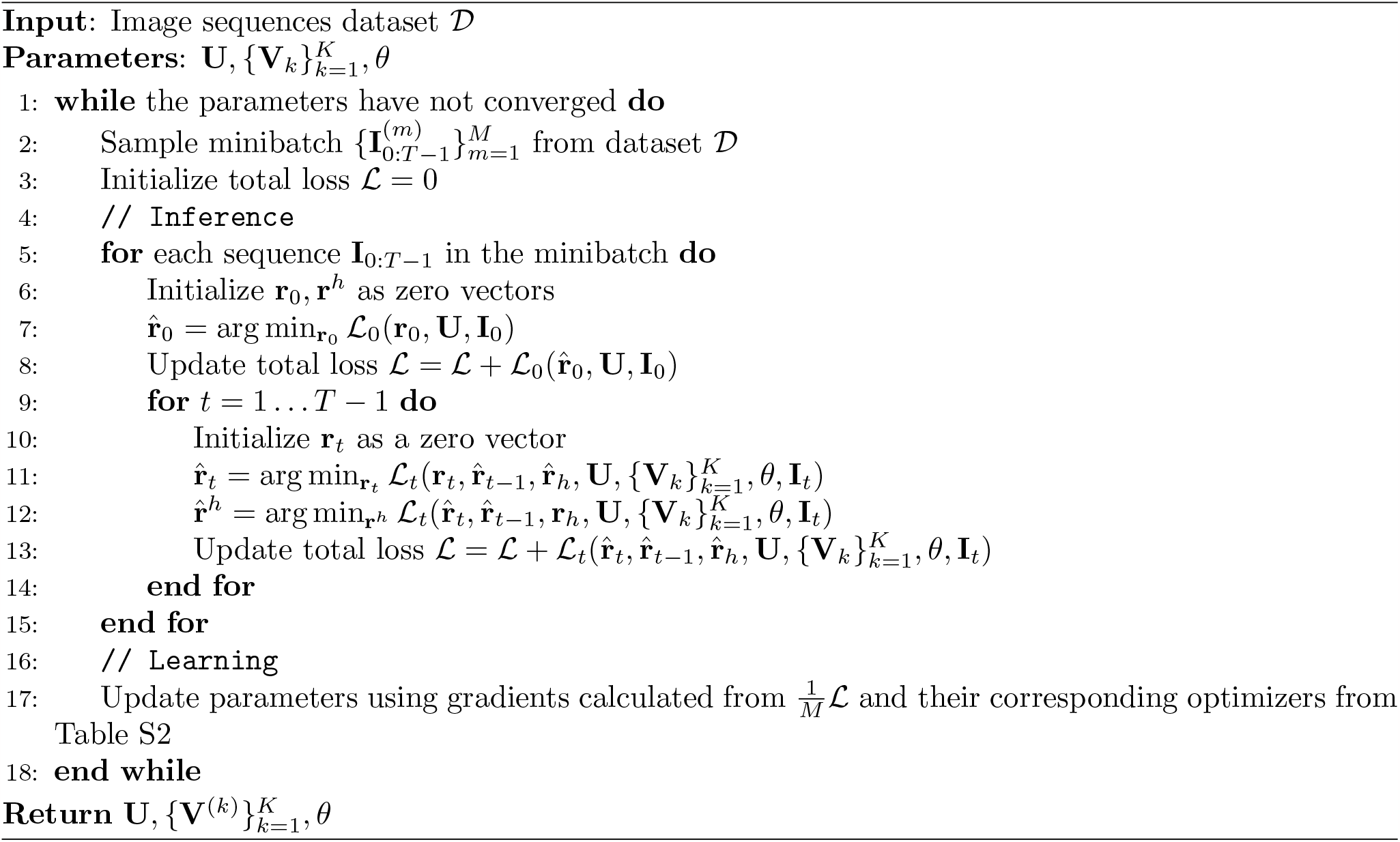

### Summary of the memory model

Table S3 summarizes the parameters and values used in the associative memory model. Table S4 summarizes the optimizers and learning rates used for inference and learning in the memory layer.

Figure S2 shows examples of cue-specific sequence recall by the associative memory model after training on different sequences. The upper left quadrant shows the image sequence used in our experiment in Figure 5, and the image sequence recalled by the trained memory-augmented DPC network. To test that the recall is cue-specific, we additionally conditioned the network with three sequences with different digits and dynamics. As the rest of Figure S2 shows, the memory-augmented DPC network successfully recalled the correct dynamics associated with the different digit cues, demonstrating the cue-specificity of the network.

### Three-level DPC model

We first describe the procedure for computing the threshold for detecting large first-level prediction errors. Using the pretrained two-level DPC model, we collect the distribution of all first-level prediction errors 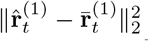 as defined in Equation 17. Figure S3(a) shows the distribution of prediction errors on the training set. We set the threshold *ρ* = 0.73 where the cumulative density of the error distribution (orange curve) reaches 0.75. To verify the efficacy of this threshold, in Figure S3(b) we show example input sequences from the test set, where the red arrows mark the time steps when the first-level prediction errors exceeded the threshold *ρ*. As shown, using this threshold robustly finds the moments when the input dynamics changed, in both straight bouncing sequences and clockwise bouncing sequences.

Table S5 describes the additional parameters and values in the three-level DPC model, in addition to those shown in Table S1 for the two-level model. The third-level top-down dynamics prediction network 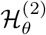 is a multi-layer perceptron that is identical to the previously described second-level network 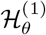, except the last output layer that has output dimension *K*^(2)^ = 2. Table S6 summarizes the optimizers and learning rates used for inference and learning in the three-level DPC experiments. Lastly, we describe the inference and learning process of the three-level model in Algorithm S2.

#### Algorithm S2 Inference & learning process for the three-level DPC model

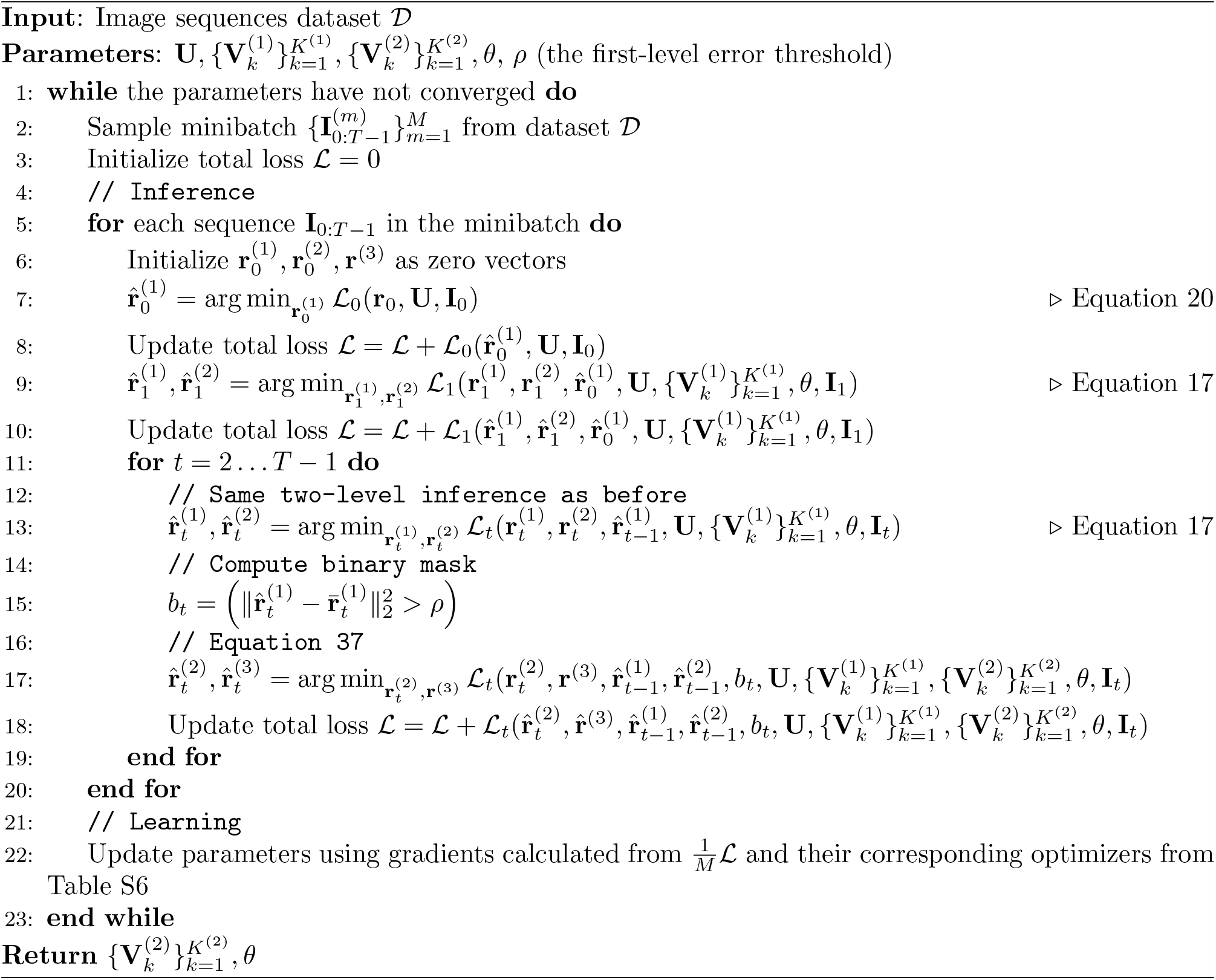

**Figure S1:**
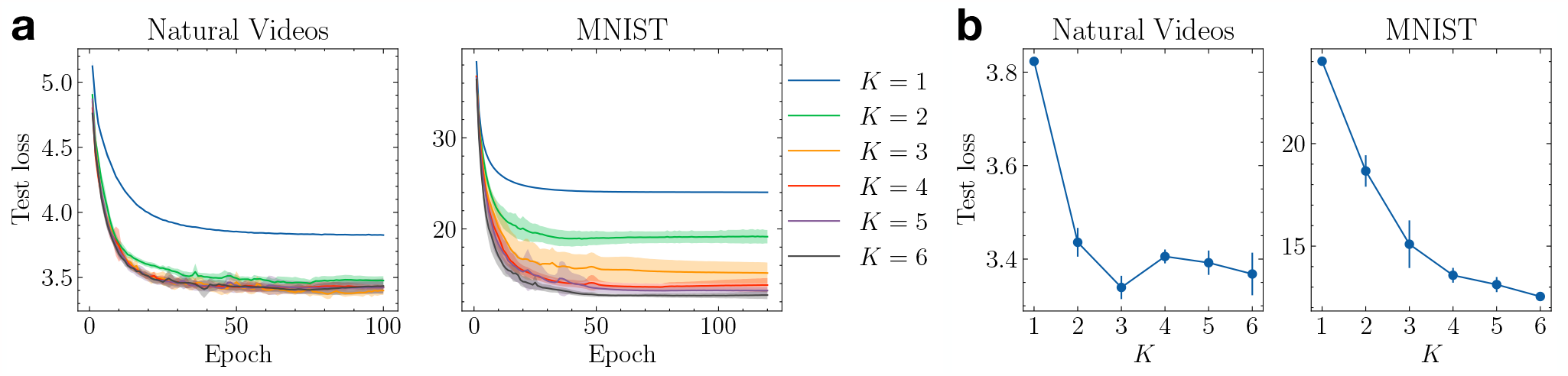
Improvement on test set loss saturates as the number of transition matrices increases. **(a)** Test set loss as training proceeded. Shaded area denotes *±* 1 standard deviation computed over eight runs with random initialization for each *K. K* = 1 shows the performance of the single-layer model. **(b)** Best test loss as *K* increases. Error bars denote *±*1 standard deviation.

**Figure S2:**
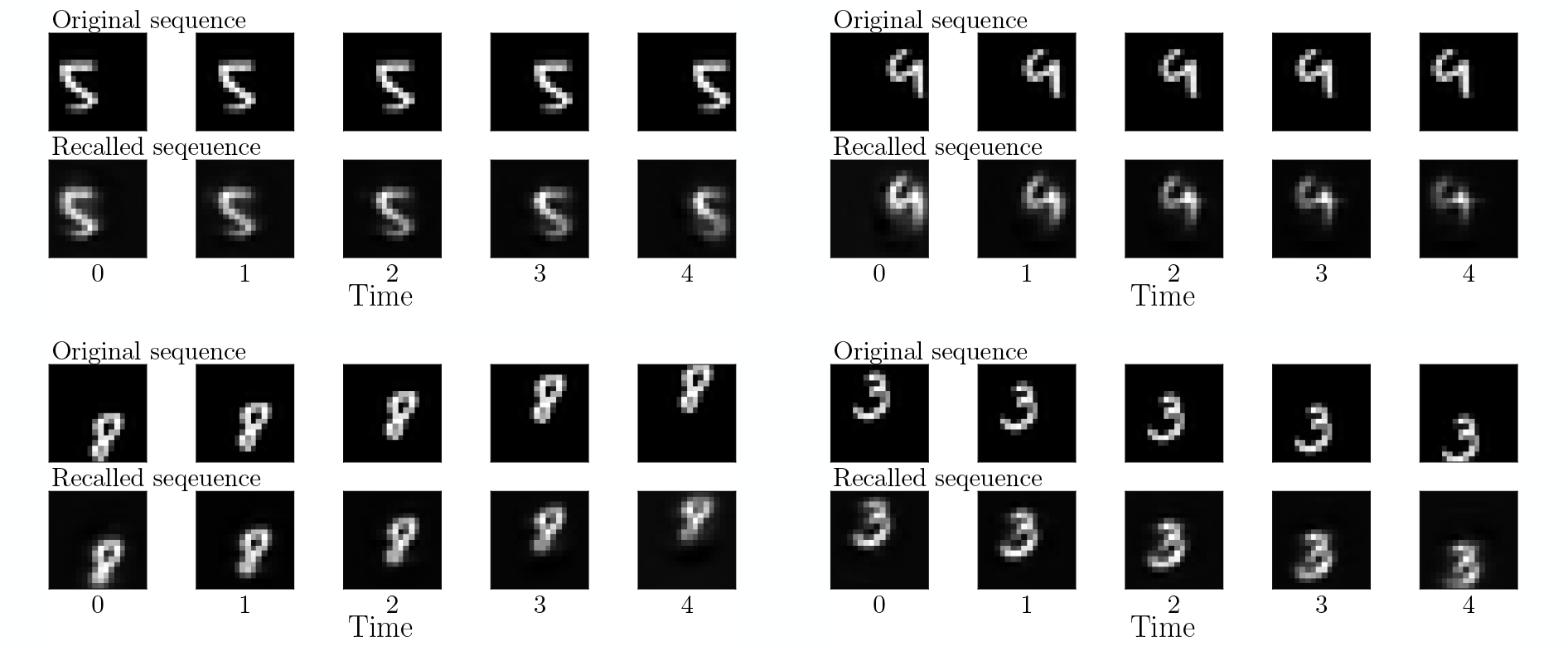
Cue-triggered recall is cue-specific. Four examples of cue-specific sequence recall by the associative memory model after training on different sequences, when given the first frame as the cue. In each quadrant: top: the original image sequence; bottom: cue-triggered recall of the stored sequence.

**Figure S3:**
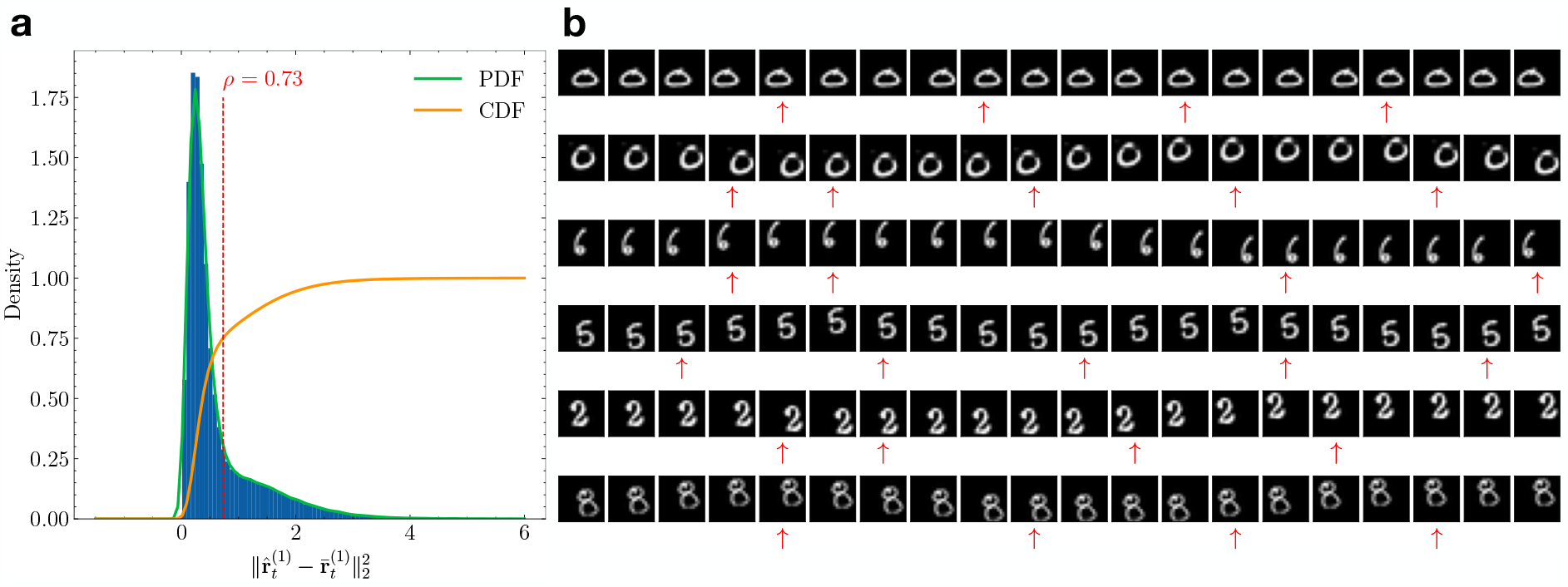
Prediction error threshold robustly finds changes of dynamics. **(a)** The distribution of first-level prediction errors in the two-level DPC model on the Moving MNIST training set. The red dashed line denotes the threshold *ρ* = 0.73, where the cumulative density reaches 0.75. **(b)** Examples of input sequences in the test set. The red arrows mark time steps when the first-level prediction errors exceeded *ρ*, corresponding to changes in input dynamics.

**Table S1:** DPC generative model parameters and values.

| Parameter | Description | Value (Natural/MNIST) |
| --- | --- | --- |
| $\mathbf{r}_t \in \mathbb{R}^N \quad \forall t = 0 \dots T-1$ | Lower-level latent variable from time 0 to $T-1$ | $N = 512/648$ |
| $\mathbf{r}^h \in \mathbb{R}^{N_h}$ | Higher-level latent variable | $N_h = 20/20$ |
| $\mathbf{I}_t \in \mathbb{R}^M \quad \forall t = 0 \dots T-1$ | Input observations from time 0 to $T-1$ | $M = 256/324$ |
| $\mathbf{U} \in \mathbb{R}^{M \times N}$ | Learnable spatial filters | Learned |
| $\sigma^2 \in \mathbb{R}$ | Spatial variance | 1 / 1 |
| $\sigma_r^2 \in \mathbb{R}$ | Temporal variance | 0.4 / 0.4 |
| $K$ | Number of transition matrices | $K = 5/5$ |
| $\mathbf{V}_k \in \mathbb{R}^{N \times N} \quad \forall k = 1 \dots K$ | Transition matrices | Learned |
| $\mathcal{H}_\theta$ | Top-down dynamics network | Learned |
| $\lambda$ | Sparsity penalty for $\mathbf{r}_t$ | $5 \times 10^{-4} / 1 \times 10^{-3}$ |
| $\lambda_h$ | Prior penalty for $\mathbf{r}^h$ | $5 \times 10^{-6} / 5 \times 10^{-6}$ |

**Table S2:** Optimizers and learning rates used for inference and learning in the DPC experiments. Here Δ denotes the difference in **r**_*t*_ or **r**^*h*^ from before and after the current iteration of gradient descent.

| Optimizee / Other Hyperparameters | Optimizer (learning rate: Natural/MNIST) / Value |
| --- | --- |
| <u>Inference</u> |  |
| $\mathbf{r}_t \quad \forall t = 0 \dots T-1$ | SGD (3/2.5) |
| $\mathbf{r}^h$ | ADAM ($1 \times 10^{-3} / 1 \times 10^{-3}$) |
| Convergence criteria | $\ \Delta\ _2 \leq 0.01$ |
| <u>Learning</u> |  |
| $\mathbf{U}$ | SGD (2.5/0.5) |
| $\mathbf{V}_k \in \mathbb{R}^{N \times N} \quad \forall k = 1 \dots K$ | ADAM ($5 \times 10^{-4} / 1 \times 10^{-3}$) |
| $\mathcal{H}_\theta$ | ADAM ($5 \times 10^{-4} / 1 \times 10^{-3}$) |
| Batch size | 512/512 |
| Epochs | 100/120 |
| Learning rate exponential decay | 0.98 / 0.985 |

**Table S3:** Memory model parameters and values.

| Parameter | Description | Value |
| --- | --- | --- |
| $\mathbf{m} \in \mathbb{R}^{N_m}$ | memory latent variable from time 1 to $T$ | $N_m = 256$ |
| $\mathbf{G} \in \mathbb{R}^{(N+N_h) \times N_m}$ | Memory weights | Learned |
| $\lambda_m$ | Prior penalty for $\mathbf{m}$ | 0 |

**Table S4:** Optimizers and learning rates used for inference and learning in the memory model. Here Δ denotes the difference in **m** from before and after the current iteration of gradient descent.

| Optimizee / Other Hyperparameters | Optimizer (learning rate) / Value |
| --- | --- |
| <u>Inference</u> |  |
| **m** | ADAM (0.005) |
| Convergence criteria | $\ \Delta\ _2 \leq 0.01$ |
| <u>Learning</u> |  |
| **G** | ADAM (0.001) |
| Epochs | 5 |
| Learning rate exponential decay | 0.99 |

**Table S5:** Additional parameters and values for the three-level DPC model.

| Parameter | Description | Value |
| --- | --- | --- |
| $\mathbf{r}^{(3)} \in \mathbb{R}^{N^{(3)}}$ | Third-level latent variable | $N^{(3)} = 10$ |
| $\sigma_{r^{(2)}}^2 \in \mathbb{R}$ | Variance of second-level prediction error | 1 |
| $K^{(2)}$ | Number of second-level transition matrices | 2 |
| $\mathbf{V}_k^{(2)} \in \mathbb{R}^{N^{(2)} \times N^{(2)}} \quad \forall k = 1 \dots K^{(2)}$ | Learnable second-level transition matrices | Learned |
| $\mathcal{H}_\theta^{(2)}$ | Third-level top-down dynamics network | Learned |

**Table S6:** Additional optimizers and learning rates used for inference and learning in the three-level DPC experiments.

| Optimizee / Criteria | Optimizer / Value |
| --- | --- |
| <u>Inference</u> |  |
| $\mathbf{r}_t^{(2)} \quad \forall t = 1 \dots T - 1$ | ADAM ($1 \times 10^{-3}$) |
| $\mathbf{r}^{(3)}$ | ADAM ($1 \times 10^{-3}$) |
| Convergence criterion | $\ \Delta\ _2 \leq 0.01$ |
| <u>Learning</u> |  |
| $\mathbf{V}_k \in \mathbb{R}^{N^{(2)} \times N^{(2)}} \quad \forall k = 1 \dots K^{(2)}$ | ADAM ($1 \times 10^{-3}$) |
| $\mathcal{H}_\theta^{(2)}$ | ADAM ($1 \times 10^{-3}$) |

